# Neuronal DNA repair reveals strategies to influence CRISPR editing outcomes

**DOI:** 10.1101/2024.06.25.600517

**Authors:** Gokul N Ramadoss, Samali J Namaganda, Jennifer R Hamilton, Rohit Sharma, Karena G Chow, Bria L Macklin, Mengyuan Sun, Jia-Cheng Liu, Christof Fellmann, Hannah L Watry, Julianne Jin, Barbara S Perez, Cindy R Sandoval Espinoza, Madeline P Matia, Serena H Lu, Luke M Judge, Andre Nussenzweig, Britt Adamson, Niren Murthy, Jennifer A Doudna, Martin Kampmann, Bruce R Conklin

**Affiliations:** Gladstone Institutes, San Francisco, CA, 94158, USA; Institute for Neurodegenerative Diseases, University of California, San Francisco, CA, 94158, USA; Innovative Genomics Institute, University of California, Berkeley, CA, 94720, USA; Department of Molecular & Cell Biology, University of California, Berkeley, CA, 94720, USA; Department of Bioengineering, University of California, Berkeley, CA, 94720, USA; Laboratory of Genome Integrity, National Cancer Institute, NIH, Bethesda, MD, 20892, USA; Department of Cellular & Molecular Pharmacology, University of California, San Francisco, CA, 94158, USA; Department of Pediatrics, University of California, San Francisco, CA, 94158, USA; Department of Molecular Biology, Princeton University, Princeton, NJ, 08544, USA; Lewis–Sigler Institute for Integrative Genomics, Princeton University, Princeton, NJ, 08544, USA; California Institute for Quantitative Biosciences, University of California, Berkeley, CA, 94720, USA; Howard Hughes Medical Institute, University of California, Berkeley, CA, 94720, USA; Department of Chemistry, University of California, Berkeley, CA, 94720, USA; MBIB Division, Lawrence Berkeley National Laboratory, Berkeley, CA, 94720, USA; Department of Biochemistry & Biophysics, University of California, San Francisco, CA, 94158, USA; Department of Medicine, University of California, San Francisco, CA, 94158, USA

## Abstract

Genome editing is poised to revolutionize treatment of genetic diseases, but poor understanding and control of DNA repair outcomes hinders its therapeutic potential. DNA repair is especially understudied in nondividing cells like neurons, which must withstand decades of DNA damage without replicating. This lack of knowledge limits the efficiency and precision of genome editing in clinically relevant cells. To address this, we used induced pluripotent stem cells (iPSCs) and iPSC-derived neurons to examine how postmitotic human neurons repair Cas9-induced DNA damage. We discovered that neurons can take weeks to fully resolve this damage, compared to just days in isogenic iPSCs. Furthermore, Cas9-treated neurons upregulated unexpected DNA repair genes, including factors canonically associated with replication. Manipulating this response with chemical or genetic perturbations allowed us to direct neuronal repair toward desired editing outcomes. By studying DNA repair in postmitotic human cells, we uncovered unforeseen challenges and opportunities for precise therapeutic editing.

## INTRODUCTION

Thousands of genetic diseases could be corrected by precise genomic edits, using tools such as CRISPR-Cas9 to induce perturbations at targeted locations in the genome^1,2^. However, a fundamental roadblock is our inability to control how those perturbations are repaired^3^. CRISPR nucleases, base editors, and prime editors perturb DNA in different ways^4–7^, but in each case, the editing outcome is ultimately determined by how the cellular DNA repair machinery responds to that perturbation^8–10^. Repair that restores the original sequence instead of editing it is unproductive, and imprecise repair can cause harmful unintended changes^3^. To ensure that the desired edit occurs in each cell, therapeutic genome editing requires thorough understanding and control of DNA repair.

Surprisingly little is known about DNA repair in postmitotic cells such as neurons, which cannot regenerate yet must withstand an entire lifetime’s worth of DNA damage. This gap in understanding hinders research into many diseases such as neurodegeneration and aging, and also limits our control over CRISPR editing outcomes. Many neurodegenerative diseases are caused by dominant genetic mutations, making them strong candidates for CRISPR-based gene inactivation^11–16^. Cas9-induced double strand breaks (DSBs) can disrupt these mutant alleles and reverse disease phenotypes. However, this requires specific DSB repair outcomes that produce the proper insertion/deletion mutations (indels) capable of frameshifting and eliminating the toxic gene product^17^.

Whether the DSB results in a desired indel or not is determined by the competing DSB repair pathways active in the cell (**Fig. 1a**, **Fig. S1**). In fact, differential expression of even a single DNA repair gene can change a cell’s editing outcome^8^. DSB repair pathways in nondividing cells likely differ drastically from those in the rapidly-proliferating and transformed cell lines used by most editing studies to date^18–21^. Pathways such as homology directed repair (HDR) for example, which are restricted to certain stages of the cell cycle, should be inactive in non-cycling cells^22^. Furthermore, DSB repair may be particularly unique in neurons, where some early-response genes are activated by the presence of DSBs in their own promoters^20^, and DSBs have even been implicated in memory formation^23^. Therefore, we hypothesized that the rules of CRISPR editing outcomes may differ in postmitotic neurons compared to the dividing cells that have shaped the literature thus far.

**Figure 1:**
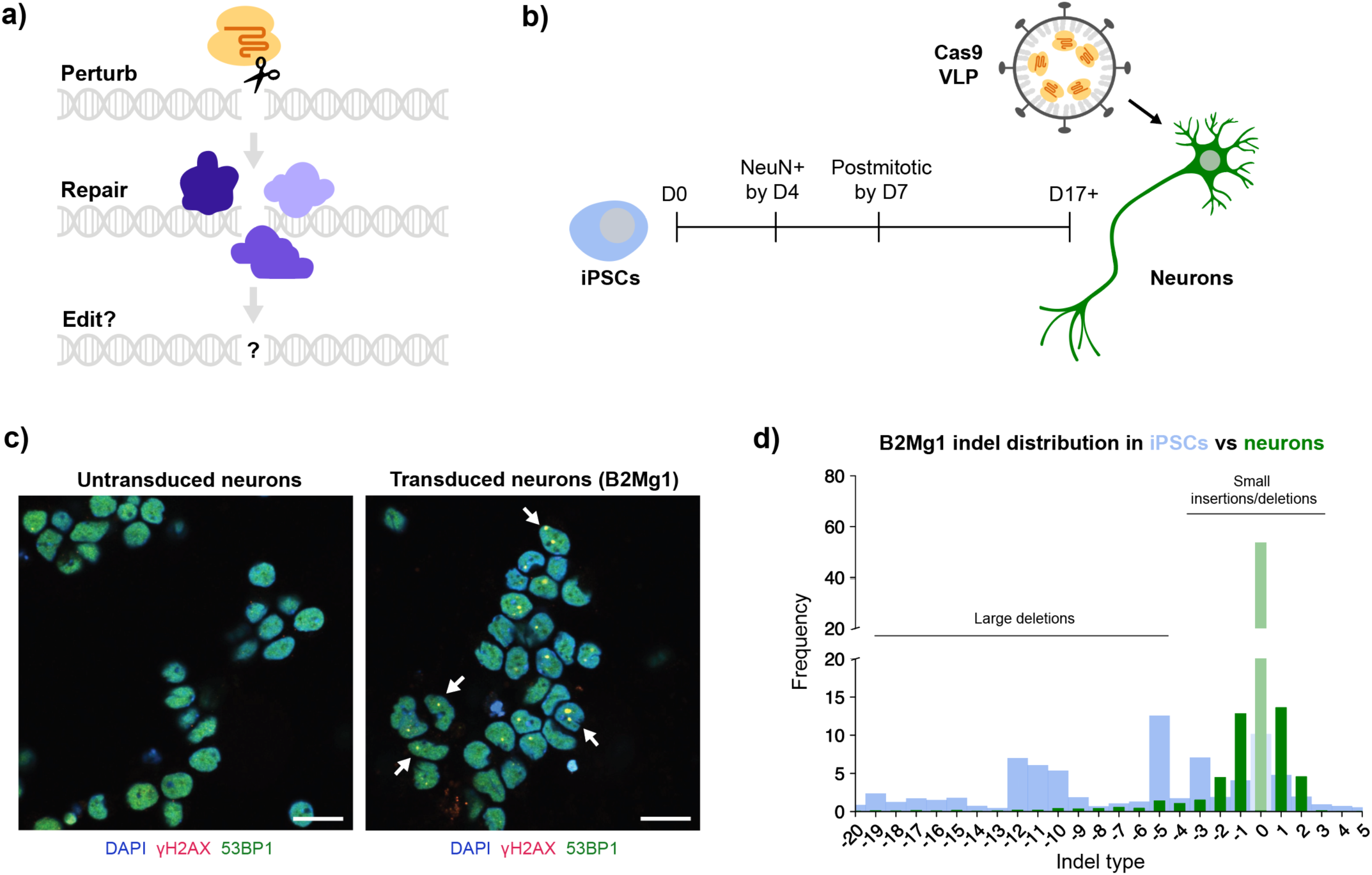
Modeling CRISPR repair outcomes in postmitotic human neurons. **a)** Schematic: Genome editing proteins can perturb DNA, but cellular DNA repair determines the editing outcome. **b)** Timeline of differentiating iPSCs (blue) into neurons (green). After at least 2 weeks of differentiation/maturation, postmitotic neurons are treated with VLPs delivering Cas9 protein (yellow) and sgRNA (orange). **c)** Cas9 VLPs induce DSBs in human iPSC-derived neurons. Representative ICC images of neurons 3 days post-transduction with B2Mg1 VLPs, and age-matched untransduced neurons. Scale bar is 20 µm. Arrows denote examples of DSB foci: yellow puncta co-labeled by γH2AX (red) and 53BP1 (green). Dose: 1 µL VLP (FMLV) per 100 µL media. **d)** Genome editing outcomes differ between iPSCs and isogenic neurons. CRISPResso2 analysis of amplicon-NGS, from cells 4 days post-transduction with B2Mg1 VLPs. Dose: 2 µL VLP (HIV) per 100 µL media. Data are averaged across 6 replicate wells per cell type transduced in parallel, and expressed as a percentage of total reads. Thick blue background bars are from iPSCs; thin green foreground bars are from neurons.

To test this hypothesis, we compared how human induced pluripotent stem cells (iPSCs) and iPSC-derived neurons respond to Cas9-induced DNA damage. We discovered that compared to these isogenic dividing cells, neurons accumulate indels over a longer time period, and upregulate unexpected DNA repair genes in response to Cas9 exposure. Furthermore, we showed that manipulating this repair response can influence the efficiency and precision of genome editing in neurons, adding important new tools to the genome modification toolkit.

## RESULTS

### Virus-like particles efficiently deliver Cas9 to human iPSC-derived neurons

To investigate how Cas9-induced DSBs are repaired in neurons, we first needed a platform to deliver controlled amounts of Cas9 into postmitotic human neurons. We used a well-characterized protocol^24,25^ to differentiate human iPSCs into cortical-like excitatory neurons (**Fig. 1b**). Immunocytochemistry (ICC) confirmed the purity of these iPSC-derived neurons. Over 99% of cells were Ki67-negative by Day 7 of differentiation, and approximately 95% of cells were NeuN-positive from Day 4 onward (**Fig. S2**). These observations confirm that within one week our cells rapidly become postmitotic, and uniformly express key neuronal markers.

While iPSCs and other dividing cells are amenable to electroporation and chemical transfection, transient Cas9 delivery to neurons remains challenging. Recently, virus-like particles (VLPs) inspired by Friend murine leukemia virus (FMLV) and human immunodeficiency virus (HIV) have been shown to successfully deliver CRISPR enzymes to many mouse tissues, including mouse brain^26–29^. Unlike viruses, which deliver genomic material into cells, VLPs are engineered to deliver protein cargo such as Cas9. Viruses pseudotyped with the glycoprotein VSVG are known to transduce LDLR-expressing cells including neurons^30^, and co-pseudotyping particles with the envelope protein BaEVRless (BRL) has been shown to improve transduction in multiple human cell types^31^. Therefore, we reasoned that VLPs pseudotyped with VSVG and/or BRL could efficiently transduce human neurons.

We produced VLPs containing Cas9 ribonucleoprotein (RNP) to induce DSBs, with or without an mNeonGreen transgene to track transduction. By flow cytometry, we found that multiple types of VLPs effectively delivered cargo to our neurons, with up to 97% efficiency (**Fig. S3**). For subsequent experiments, we proceeded with two particles interchangeably: VSVG pseudotyped HIV VLPs (also known as enveloped delivery vehicles^27^), or VSVG/BRL co-pseudotyped FMLV VLPs. Furthermore, ICC confirmed that Cas9-VLPs successfully induced DSBs in our neurons, co-labeled by markers gamma-H2AX (γH2AX) and 53BP1 (**Fig. 1c**, **Fig. S4**). This platform to acutely perturb DNA in human neurons enables the study of DNA repair in clinically relevant postmitotic cells.

### CRISPR repair outcomes differ in neurons compared to dividing cells

To examine how neurons repair DSBs, we used VLPs to deliver identical doses of Cas9 RNP into human iPSC-derived neurons and isogenic iPSCs. We selected a single-guide RNA (sgRNA), B2Mg1, that yields a variety of indel types in iPSCs, suggesting it is compatible with multiple DSB repair pathways. End resection-dependent DSB repair pathways such as microhomology-mediated end joining (MMEJ) are typically restricted to certain stages of the cell cycle (S/G2/M), while nonhomologous end joining (NHEJ) is not^22,32,33^. Since postmitotic cells have exited the cell cycle, they are predicted to predominantly utilize NHEJ when repairing DSBs.

Indeed, while B2Mg1-edited iPSCs displayed a broad range of indels, neurons exhibited a much narrower distribution of outcomes (**Fig. 1d**). In iPSCs, the most prevalent indel outcomes were larger deletions typically associated with MMEJ, as expected for dividing cells^33^. In neurons, the most prevalent outcomes were those usually attributed to NHEJ: small indels associated with NHEJ processing, and unedited outcomes caused by either indel-free classical NHEJ (cNHEJ) or lack of Cas9 cutting^34,35^. This was true for several different sgRNAs tested. Even though each sgRNA had a different intrinsic distribution of available indel types, in each case, the MMEJ-like larger deletions were predominant in iPSCs, and the NHEJ-like smaller indels were predominant in neurons (**Fig. S5**). These results suggest that postmitotic neurons employ different DSB repair pathways than isogenic dividing cells, yielding different CRISPR editing outcomes.

Unresolved DSBs can be lethal to cycling cells, as DNA damage checkpoints trigger cell cycle arrest and/or apoptosis^36,37^. Therefore, for dividing cells, resolving a DSB mutagenically can be less harmful than leaving it unrepaired. For example, mitotic cells often utilize extremely indel-prone MMEJ repair to avoid progressing through M phase with unresolved DSBs^32,33^. This is consistent with our observed editing outcomes in iPSCs. On the other hand, postmitotic cells do not face replication checkpoints, and thus might not be subjected to the same pressures. Therefore, we hypothesized that DSBs could be resolved over a longer time scale in postmitotic cells.

### Cas9-induced indels accumulate slowly in neurons

In dividing cells, the repair half-life of Cas9-induced DSBs is reportedly between 1-10 hours; even in the slowest-repaired cut sites, the fraction of unresolved DSBs peaks within just over one day^38^. DSB repair in our iPSCs matched this expected timing, with indels plateauing within a few days. In contrast, indels in neurons continued to increase for up to two weeks post-transduction (**Fig. 2a**, **Fig. S6a**).

**Figure 2:**
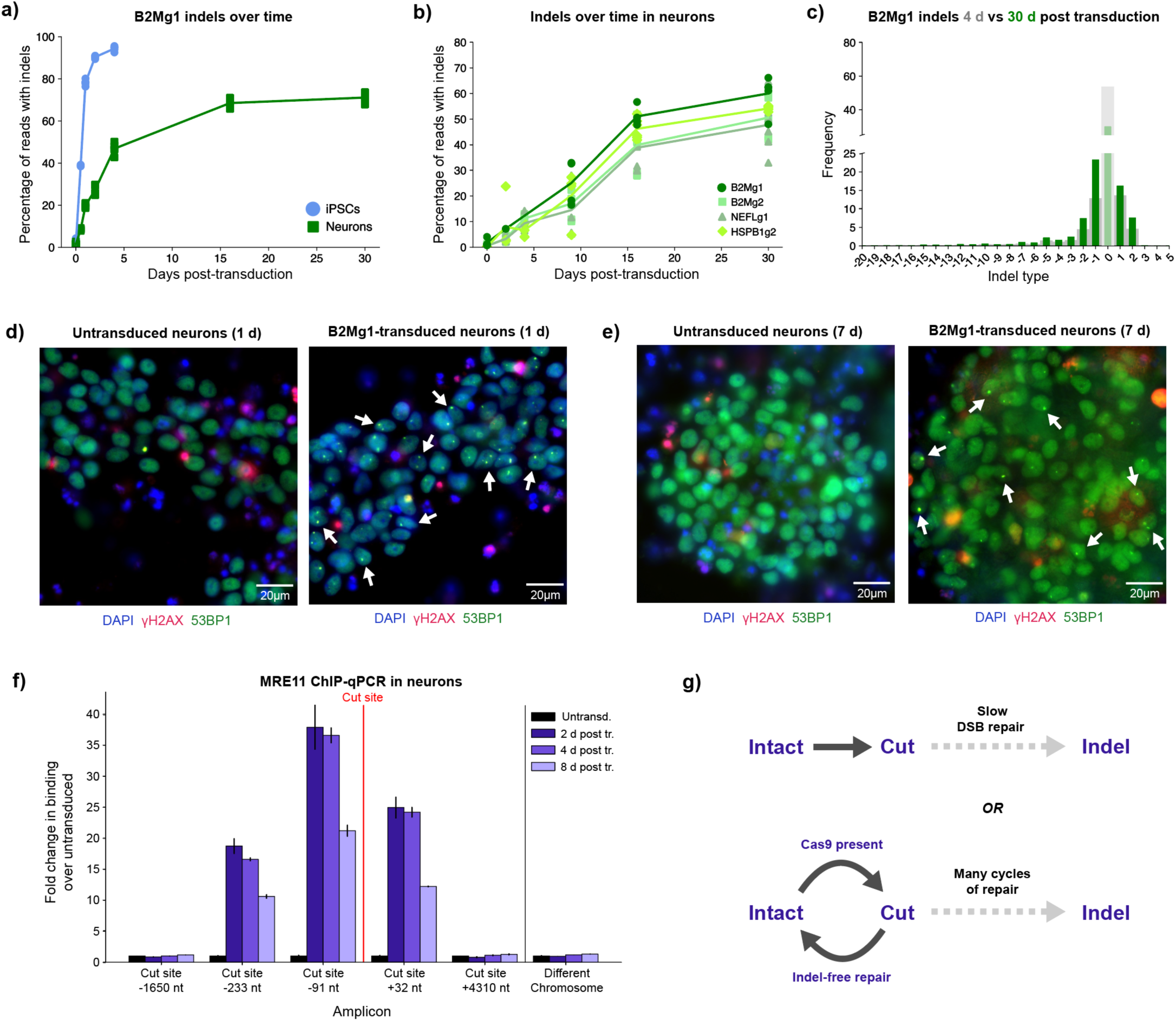
Cas9-induced indels accumulate over a prolonged time span in neurons. **a)** Cas9-induced indels accumulate more slowly in neurons than in iPSCs. Dose: 2 µL B2Mg1 VLP (HIV) per 100 µL media. For a-b: points represent individual replicates (some obscured by overlap); curves connect means at each timepoint. For a-c: 6 replicate wells per condition transduced in parallel. CRISPResso2 analysis of amplicon-NGS. **b)** Several sgRNAs show weeks-long accumulation of indels in neurons. Dose: 1 µL VLP (FMLV) per 100 µL media. **c)** Insertions and deletions both increase over time in neurons. Dose: 2 µL B2Mg1 VLP (HIV) per 100 µL media. Histogram: thick gray background bars are from 4 d timepoint, and thin green foreground bars are from 30 d. **d-e)** Cas9-induced DSBs remain detectable in neurons at least 7 days post-transduction. Representative ICC images of neurons 1 day (d) and 7 days (e) post-transduction with B2Mg1 VLPs, and age-matched untransduced neurons. Dose: 2 µL B2Mg1 VLP (FMLV) per 100 µL media. Arrows denote examples of DSB foci. See Extended Data Figure S8 for unmerged/uncropped panels. **f)** MRE11 is bound near the cut site in neurons for at least 8 days post-transduction. Dose: 2 µL B2Mg1 VLP (FMLV) per 100 µL media. Binding events quantified by ChIP-qPCR for each amplicon, normalized for amplification efficiency and input chromatin. Average of 3 replicate ChIP-qPCR reactions, normalized to untransduced control for each amplicon. Error bars show SD. **g)** Schematic of two possible models for prolonged indel accumulation in neurons.

We tested multiple sgRNAs including disease-relevant targets. Surprisingly, for every sgRNA, neuron indels continued to increase for at least 16 days post-delivery of transient Cas9 RNP (**Fig. 2b**). Regardless of the intrinsic indel distribution, each available indel type for each sgRNA increased in frequency for weeks (**Fig. 2c**, **Fig. S6b-d**). Additionally, this extended time course of editing was replicated by both types of VLPs (**Fig. S6e-h**).

We found no evidence that this prolonged indel accumulation in neurons was influenced by proliferating cells (**Fig. S2**), inefficient transduction (**Fig. S3**), or residual VLP in the media (**Fig. S7a**). Furthermore, to test whether this phenomenon was specific to DSB repair, we transduced neurons and iPSCs with VLPs delivering an adenine base editor (ABE) instead of Cas9. Using the same delivery particle but engaging a different DNA repair pathway than DSBs, ABE-VLP-mediated editing in neurons was comparably efficient to iPSCs – and sometimes even more efficient – even within only three days post transduction (**Fig. S7b**). This result suggests that the slow indel accumulation is DSB repair-specific, and not caused by deficits in neuronal VLP delivery.

Interestingly, we observed a similarly slow timeline of indel accumulation in postmitotic iPSC-derived cardiomyocytes (**Fig. S7c**). Therefore, this prolonged indel accumulation might also apply to other clinically relevant postmitotic cells, not only neurons.

This weeks-long timeline of editing could have major clinical implications. Gene inactivation therapies in nondividing tissues might take longer than anticipated to be effective, and both on-target and off-target editing may accumulate over longer intervals. Additionally, persistent DSBs in neurons have been associated with genomic instability and even neurodegeneration^39–41^, so characterizing the duration of Cas9-induced damage and repair is critical.

### DSB repair is detectable in neurons for more than one week post Cas9 delivery

To assess the duration of this damage in neurons, we measured multiple signals of DSB repair over time after delivering transient Cas9 RNP via VLPs. DSB foci (γH2AX/53BP1) were strongly detectable by ICC as early as one day post-transduction, confirming efficient delivery and rapid induction of DSBs in neurons (**Fig. 2d**). Interestingly, DSB foci remained detectable in neurons for at least seven days post-transduction (**Fig. 2e**). Persistent DSB repair signal was observed for sgRNAs targeting both lowly-transcribed (*B2M*) and highly-transcribed (*NEFL*) genes (**Fig. S8**). This long-lived repair signal is consistent with the prolonged accumulation of indels in neurons. DSB foci in iPSCs cannot be compared over the same span, as proliferating cells replicate many times within a week, and any unresolved signal would be diluted.

To more quantitatively measure this repair in neurons, we used chromatin immunoprecipitation with quantitative real-time PCR (ChIP-qPCR) to measure the binding of repair proteins Mre11 and γH2AX near the cut site, at several timepoints post-transduction. Mre11 binding was strongly detected within a few hundred bases of the cut site, and only in transduced samples (**Fig. 2f**), matching patterns seen in other cell types^42^. But intriguingly, Mre11 binding near the cut site remained strongly detected in neurons even 8 days post-transduction, decreasing by only ∼50% between days 2 and 8.

As expected based on previous reports^42,43^, γH2AX binding was much broader, with maximal signal detected several kilobases away from the cut site. Interestingly, while γH2AX binding >100 bases from the cut site decreased to background levels between days 2 and 8, γH2AX binding near the cut site only decreased by ∼50% during this interval (**Fig. S9a**).

Some binding at each timepoint can be attributed to DSBs that had already been resealed, as illustrated by an amplicon that spans across the cut site and thus should only amplify if the cut was resealed (**Fig. S9b-d**). Cut sites resealed without an indel can be repeatedly recut by any remaining Cas9 RNP, until an indel prevents subsequent Cas9 binding. This is consistent with the slow indel accumulation observed in neurons. These results cannot be compared to dividing cells like iPSCs, where one locus would become many due to replication, and Cas9 protein would get rapidly diluted within 2 days (**Fig. S9e**).

Altogether, these findings suggest that postmitotic neurons either take longer to complete DSB repair, or undergo more cycles of indel-free repair and recutting until indels arise, or perhaps both (**Fig. 2g**). Either way, our results confirm that DSB repair signals at the target site persisted in neurons for more than one week post-delivery of Cas9 RNP – much longer than expected.

### Cas9-VLPs elicit a striking transcription-level response in neurons

Based on this unexpectedly prolonged time scale of editing, we hypothesized that neuronal DNA repair could include transcription-level regulation, not only post-translational regulation. To test this, we used bulk RNA sequencing (RNA-seq) to characterize differentially expressed genes (DEGs) in iPSCs and neurons transduced with Cas9-VLP, relative to untransduced cells (**Supplemental Table 1**). Unlike transduced iPSCs, transduced neurons exhibited a skewed transcriptional response, with far more genes upregulated than downregulated (**Fig. 3a-b**). This neuron-specific response was replicated with 3 different sgRNA conditions: *B2M*-targeting (B2Mg1), *NEFL*-targeting (NEFLg1), or non-targeting (NTg1) sgRNAs (**Fig. S10a-d**). For every VLP condition, less than 10% of neuron DEGs were shared with iPSCs (**Fig. 3c**), indicating a distinct difference between the cell types’ responses.

**Figure 3:**
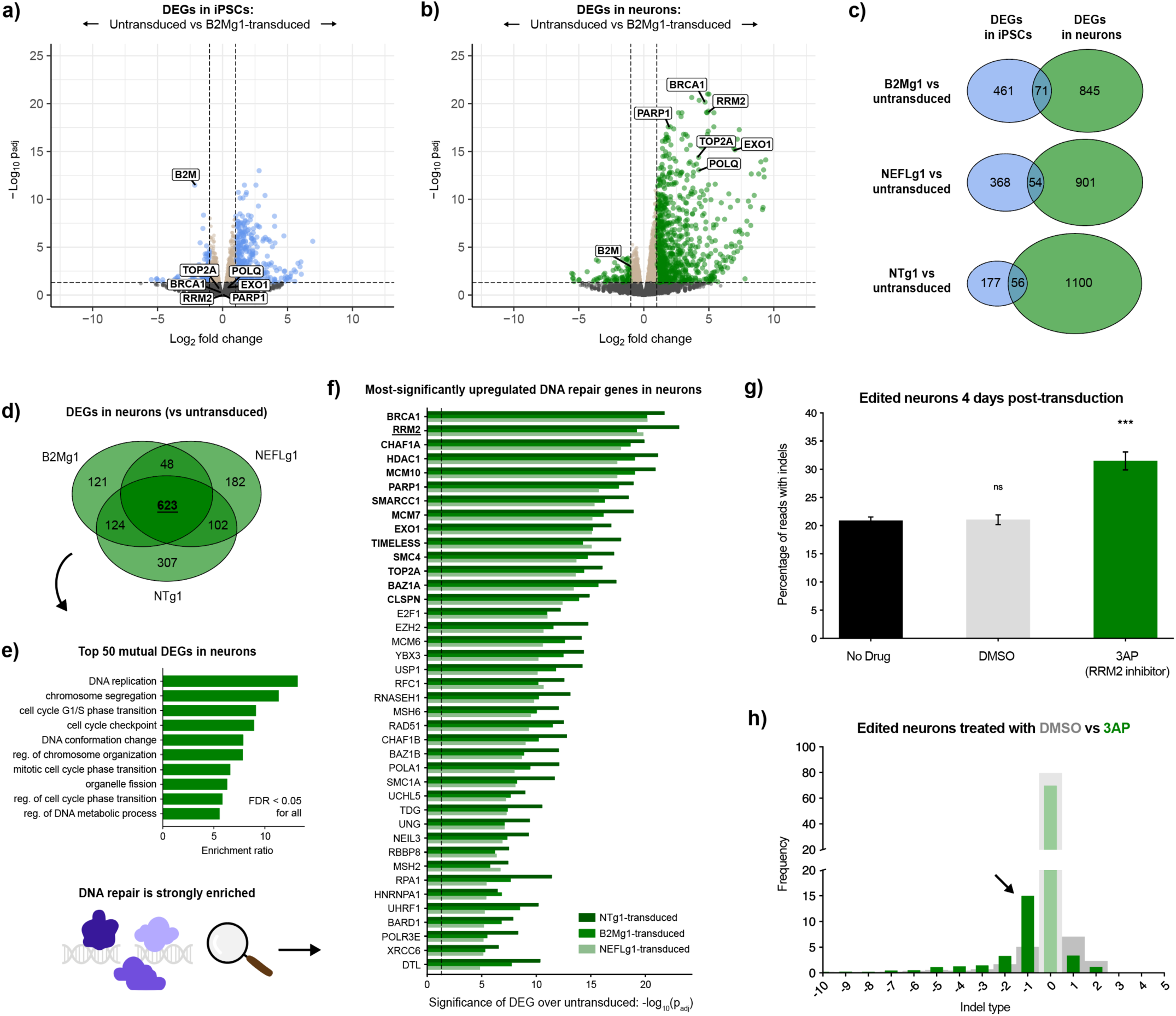
Neuronal response to Cas9-VLP reveals unexpected factors that influence editing outcomes. **a-b)** Neurons (b), but not iPSCs (a), dramatically upregulate transcription of DNA repair factors upon Cas9-VLP transduction. Volcano plots show differential expression transcriptome-wide in transduced cells relative to untransduced. Dashed lines show cutoffs for significance (padj<0.05) and effect size (fold-change >2 or <0.5). For a-f: differential expression was calculated from bulk RNA-seq across 3 replicate samples per condition transduced in parallel. Methods section details the statistical tests used to define significant DEGs. Dose: 1 µL HIV VLP per 20,000 cells (1.25 µL VLP per 100 µL media). **c)** Transduced neurons consistently have more DEGs than transduced iPSCs for 3 different sgRNAs, and <10% of DEGs are shared between the cell types. **d)** Over 75% of the DEGs in either B2Mg1- or NEFLg1-transduced neurons are shared with NTg1-transduced neurons. **e)** The most significantly altered DEGs in transduced neurons are highly enriched for DNA repair factors. **f)** Transduced neurons significantly upregulate many DNA repair genes, including factors canonically associated with replication. Top 40 DNA repair DEGs are shown, rank-ordered by averaging the adjusted p-values from each transduced condition. Bold denotes repair genes ranked in the top 50 DEGs genome-wide. **g)** Inhibiting RRM2 yields a 50% increase in neuron editing efficiency, within 4 days post-transduction. Error bars show SEM. One-Factor ANOVA with Tukey’s multiple comparison test. Each condition vs No Drug, *** p<0.0005, ns = not significant. For g-h: CRISPResso2 analysis of amplicon-NGS, averaged across 6 wells per condition transduced in parallel. Dose: 1 µL B2Mg1 VLP (FMLV) per 20,000 cells in 100 µL media. **h)** RRM2 inhibition tripled the frequency of 1-base deletions at 4 days post-transduction. Thick gray bars are DMSO condition; thin green bars are 3AP.

The neuronal response to Cas9-VLPs was remarkably consistent regardless of the sgRNA target. In fact, only two genes were differentially expressed between B2Mg1-edited and NEFLg1-edited neurons: *B2M* and *NEFL*, respectively (**Fig. S10e**). This confirms that the observed response is not locus-specific, and is not driven by loss-of-function of either targeted gene. Surprisingly, over 75% of the DEGs in B2Mg1-edited or NEFLg1-edited neurons, relative to untransduced, were also shared with NTg1-treated neurons (**Fig. 3d**, **Fig. S10f**). The top 50 DEGs shared by all three transduced neuron conditions were highly enriched for DNA repair genes (**Fig. 3e**). This suggests that Cas9-VLP induces a strong transcription-level DNA repair response in neurons, some part of which may even be DSB-independent.

### Transduced neurons upregulate unexpected DNA repair genes

The most-significantly upregulated repair genes included many pathways thought to be inactive in nondividing cells, such as end resection-related pathways^22^ (**Fig. 3f**). They also included factors known to influence prime editing and base editing^9,10^, suggesting this neuronal response could impact multiple types of editing. Additionally, transduced neurons significantly upregulated factors that respond to R-loops, single-stranded DNA, and topological stresses (**Fig. 3f**). This might explain why even NTg1-Cas9 induced a strong response: even if it does not cut, Cas9 still disrupts DNA: unwinding it, creating R-loops, and exposing single-stranded DNA^44,45^. Notably, this response was unique to neurons; DEGs were not enriched for DNA repair in any of the three transduced iPSC conditions (**Fig. S11**). These DNA repair genes were already expressed at baseline in untransduced iPSCs, whereas neurons only induced their expression upon Cas9-VLP transduction.

Intriguingly, the most-upregulated genes in transduced neurons were particularly enriched for replication-related factors, such as cell cycle checkpoints and DNA synthesis during S phase (**Fig. 3e-f**). Neurons have long been postulated to partially re-enter cell cycle following DNA damage through a process called endocycling, which replicates DNA without necessarily completing mitosis^46–49^. Cas9-VLPs could have induced such a response in our neurons. It is also possible, however, that these repair factors are canonically annotated as replication-related because they have mostly been studied in dividing cells, where their role in repairing replication-induced damage eclipses any others. In nondividing cells, these factors’ roles in responding to other types of DNA damage might be more visible. We investigated one of the strongest and most unexpected of these hits: *RRM2*.

### Transduced neurons non-canonically upregulate a subunit of ribonucleotide reductase

*RRM2* was one of the most-significantly upregulated repair genes transcriptome-wide in every transduced neuron condition (**Fig. 3f**). *RRM2* encodes a subunit of ribonucleotide reductase (RNR), the enzyme that produces deoxyribonucleoside triphosphates (dNTPs). RNR is functional when the catalytic subunit RRM1 binds one of two tightly regulated smaller subunits: RRM2 or RRM2B^50^. *RRM2* expression is canonically restricted to S phase to produce dNTPs for replication, while *RRM2B* is canonically upregulated by p53 upon DNA damage to facilitate repair^51^.

In iPSCs, each RNR subunit responded as expected: *RRM2B* was upregulated by the VLPs that induced DSBs, and *RRM2* was unaltered (**Fig. S12a-c**). In contrast, the response in neurons was completely unexpected. *RRM2B* expression was not altered in any condition. Instead, the canonically S-phase-restricted *RRM2* was one of the most upregulated genes transcriptome-wide, in every transduced neuron condition (**Fig. S12d-f**).

We reasoned that this unexpected shift in the DNA repair landscape could impact CRISPR editing. For example, in NHEJ processing where polymerase filling-in competes with other pathways^34^ (**Fig. S1**), this non-canonical RNR activation could bias the outcome by increasing nucleotide availability.

### Inhibiting these repair factors influences editing outcomes in neurons

Based on these results, we tested whether inhibiting RNR affected Cas9 editing outcomes. We treated neurons with triapine (3AP), a small molecule inhibitor of RRM2^52–55^, while delivering Cas9-VLPs targeting B2Mg1. Excitingly, 3AP treatment led to a ∼50% increase in total indels, at only four days post-transduction (**Fig. 3g**). This increase in indels came almost exclusively from boosting deletions, at the expense of insertions and indel-free repair (**Fig. 3h**). In fact, 3AP co-treatment led to a ∼3-fold increase in single-base deletions specifically, tilting the distribution toward one predictable outcome.

Two other RNR-inhibiting drugs had the same effect as 3AP on B2Mg1 editing in neurons: GW8510 which also inhibits RRM2^53,56,57^, and gemcitabine which inhibits its obligate binding partner RRM1^58,59^. Both drugs increased total indel frequency, and preferentially boosted deletions, affecting both the efficiency and precision of gene inactivation (**Fig. S13**). 3AP and gemcitabine have already been used in clinical trials for other applications^60–63^. Depending on the toxicity to dividing cells, future studies could explore their clinical relevance for enhancing therapeutic editing outcomes.

Expecting that different cut sites may differ in scission profiles and repair dependencies^64–66^, we also tested the effect of RNR inhibition on three other sgRNAs. Inhibiting RNR increased indels for B2Mg2 and NEFLg1, though without the same selectivity for single-base deletions (**Fig. S14a-d**). For HSPB1g2, RNR inhibition in fact *decreased* indels (**Fig. S14e-f**). This is consistent with the intrinsic indel distribution of HSPB1g2, which appears impermissible to the deletions that 3AP often boosts. Therefore, inhibiting RNR cannot be generalized as a method to increase indel efficiency for *all* sgRNAs. Rather, RNR inhibition influences editing outcomes in an sgRNA-dependent manner.

Overall, deciphering how clinically relevant cells respond to Cas9 unveiled unexpected DNA repair factors that influence editing outcomes. Identifying these upregulated genes highlighted many potential targets for manipulating repair. Our RNR results demonstrate that modulating these factors can reveal which outcomes they affect, and can help optimize the editing outcome for a given sgRNA of interest.

### Prolonged editing window allows manipulation of repair factors at RNA level

When modulating DNA repair factors to optimize editing outcomes, a major barrier is that not all factors are druggable. For example, in the NHEJ pathway alone, two-thirds of the factors^35^ do not have reliable small molecule inhibitors for protein-level targeting (**Fig. S15**). Since neurons activated DNA repair factors at the *transcriptional* level as well, and had a long window of days or weeks for completing repair, we reasoned that manipulating repair factors at the RNA level – rather than the protein level -- may also be sufficient to influence neuron indels. If true, this would enable modulation of any DNA repair factor, not only the druggable ones.

To test this idea, we used short interfering RNAs (siRNAs) inhibiting DNA repair genes of interest, co-encapsulated with Cas9 mRNA and sgRNA inside lipid nanoparticles (LNPs) – a delivery vehicle well-suited for all-RNA cargo (**Fig. 4a**, **Fig. S16a-c**). At baseline, LNP delivery of Cas9 and sgRNA to neurons induced a slow accumulation of indels over several days, though indel counts remained lower than with VLPs, plateauing earlier (**Fig. 4b**). To evaluate whether RNA-level inhibition could influence indel outcomes, we first targeted DNA-PKcs, a key NHEJ factor. *PRKDC*-targeting siRNAs reduced the frequency of neuron indels broadly, phenocopying small molecule inhibition of DNA-PKcs (**Fig. S16d-h**).

**Figure 4:**
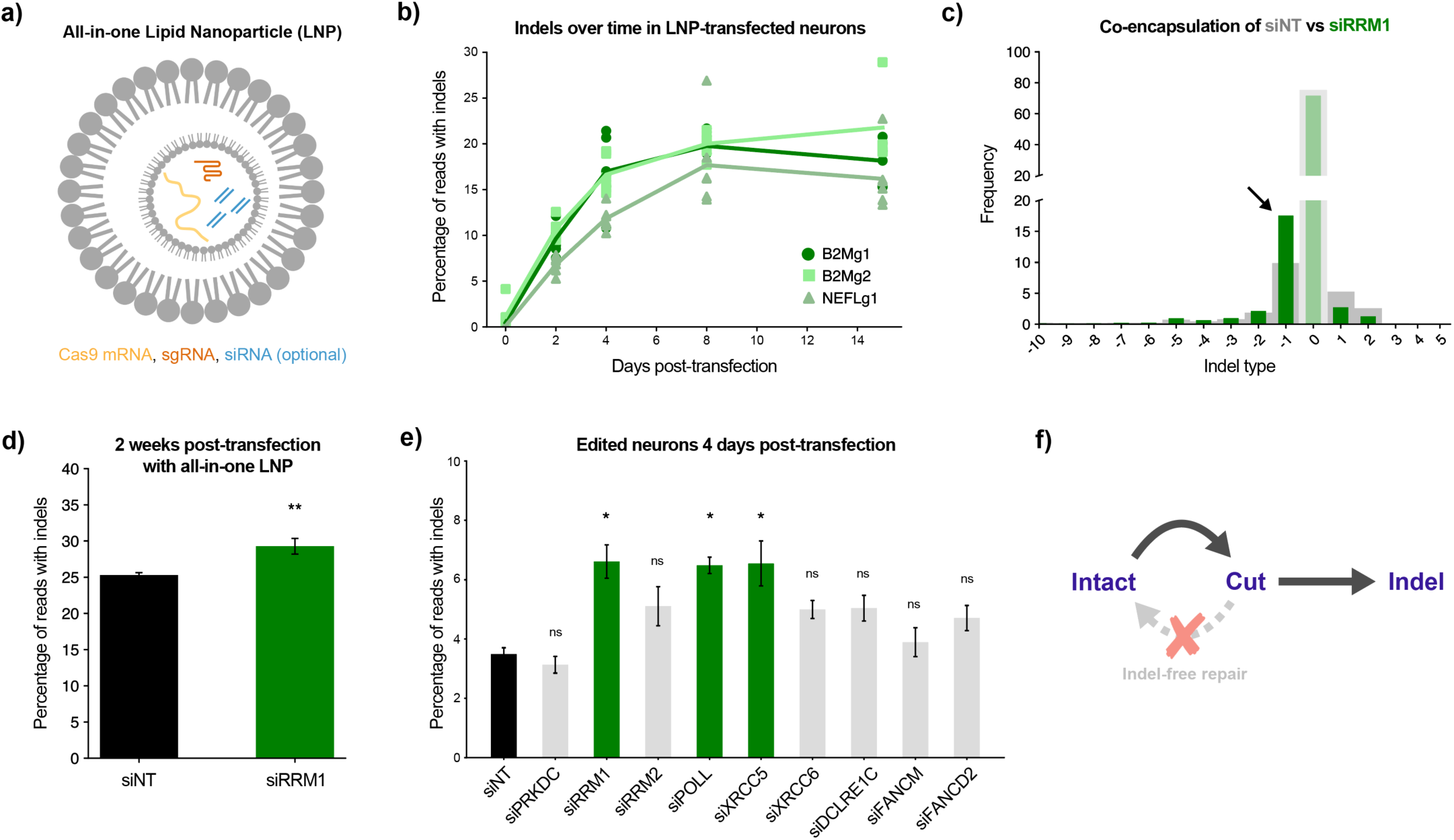
All-in-one particles deliver Cas9 and sgRNA while simultaneously manipulating DNA repair factors. **a)** Schematic illustrating all-in-one LNPs that encapsulate Cas9 mRNA (yellow) and sgRNA (orange), along with siRNAs (blue) against a repair gene of interest. **b)** Multiple sgRNAs show days-long accumulation of neuron indels following LNP transfection. Individual points represent 6 replicate wells per condition transfected in parallel (some obscured by overlap). Curves connect means at each timepoint. For b-e: CRISPResso2 analysis of amplicon-NGS from neurons. **c)** RNA inhibition of RRM1 during editing phenocopies small molecule inhibition of RRM1/2, shifting B2Mg1 editing outcomes toward deletions. Two weeks post-transfection. Thick gray background bars are from siNT condition; thin green foreground bars are from siRRM1. For c-d: averaged across 6 replicate wells per condition transfected in parallel. **d)** Co-encapsulating siRRM1 increases total B2Mg1 editing at two weeks post-transfection. Error bars show SEM. One-Factor ANOVA, ** p<0.005. **e)** All-in-one LNPs reveal additional targets that increase B2Mg1 editing efficiency at 4 days post-transfection. Averaged across 8 replicate wells per condition, transfected in parallel. Error bars show SEM. One-Factor ANOVA with Tukey’s multiple comparison test. Each condition vs siNT, * p<0.05, ns = not significant. **f)** A model for why the hits from e accelerated editing in neurons. Inhibiting indel-free repair may have directed outcomes toward indels instead of repeatedly resealing and recutting.

Next, we tested whether siRNA knockdown of RNR subunits could phenocopy small molecule inhibition. Since *RRM2* is not expressed in neurons until after Cas9 exposure – yet the siRNA is active before the Cas9 mRNA gets translated – we inhibited its obligate binding partner *RRM1*, which is more highly expressed at baseline. This siRNA treatment during B2Mg1 editing phenocopied small molecule inhibition of *RRM1/2*. Two weeks post-transfection, siRNA inhibition of *RRM1* increased the frequency of single-base deletions by ∼75%, and increased total indels by ∼20% overall (**Fig. 4c-d**). Therefore, these all-in-one LNPs allowed us to deliver editing reagents to neurons while simultaneously influencing the repair outcome with RNA interference (RNAi). This co-packaging strategy might be safer than systemically delivering drugs that are toxic to dividing cells, and it also allows us to target repair factors even if they are not druggable.

### All-in-one particles enable screening for additional repair targets that accelerate editing

To demonstrate using these tools to optimize editing, we transfected neurons with all-in-one particles targeting a small set of additional DSB repair factors – several of which are not reliably targetable by small molecule drugs. Aiming to identify perturbations that accelerate editing, we assessed editing at an earlier time point of 4 days post-transfection, before indels had plateaued. Knockdowns of *RRM1*, *POLL*, and *XRCC5* significantly increased total B2Mg1 indels by ∼80% relative to non-targeting siRNA (**Fig. 4e**). *POLL* encodes the polymerase that likely performs filling-in synthesis during NHEJ processing^34^, using the dNTPs produced by RNR. And *XRCC5* (Ku80) is one of the key factors involved in end protection to promote indel-free cNHEJ^34,35^. Our model proposes that these interventions disrupted indel-free repair of the B2Mg1 cut site, and directed the repair outcome toward indels instead (**Fig. 4f**), thus accelerating gene inactivation.

Such strategies for generating more indels at earlier timepoints could help minimize the danger of persistent DSBs. Additionally, controlling which repair pathways are utilized would improve the precision and predictability of genome editing. This platform could be repurposed to find optimal repair modifications for any sgRNA of interest, simply by encapsulating different sgRNAs and siRNAs inside the all-in-one particles.

## DISCUSSION

Altogether, our results emphasize the importance of studying genome editing therapies in the appropriate cell type models. Neurons’ distinct response to Cas9-induced DNA damage led to dramatically different repair outcomes and weeks-long accumulation of edits, which could impact both the safety and efficacy of therapeutic editing.

Investigating this response revealed that postmitotic neurons begin to express many DNA repair genes only *after* acute damage occurs – including unexpected factors like RNR which are canonically associated with cell replication. Therefore, expression levels of repair genes in *unperturbed* cells should not be used as a proxy for which repair pathways are accessible. By inhibiting RNR, we boosted the frequency of desirable indel outcomes in neurons: namely single-base deletions that should reliably lead to gene knockout. We replicated this deletion-boosting effect using four different methods of RNR inhibition including siRNA knockdown and three small molecule inhibitors – as well as two different modes of Cas9 delivery in VLPs and LNPs. Importantly, this effect was sgRNA-dependent, as different cut sites responded differently to the same repair modification.

These insights helped turn neurons’ slow indel accumulation from a challenge into an opportunity. Since the neuronal repair response was detectable for many days and involved transcription-level upregulation, we created all-in-one particles that deliver Cas9 while simultaneously manipulating the repair process via RNAi. Compared to drug inhibition, this strategy greatly expands how many factors we can target. As a proof-of-concept, we used this all-in-one screening platform to find repair modifications that accelerate indels for a given sgRNA of interest. Overall, by studying how nondividing cells repair Cas9-induced DNA damage, we discovered multiple new strategies to influence genome editing outcomes.

Several key strengths of our experimental approach enabled these findings. First, using iPSCs instead of transformed cell lines allowed us to model DNA repair in karyotypically normal cells. Second, comparing neurons to their isogenic iPSCs allowed us to evaluate CRISPR editing in dividing vs nondividing cells without confounding factors such as genetic background. Third, since iPSC-derived neurons share the genotypes and even some phenotypes of the human donors, this same platform could be used to test and optimize a genome editing therapy in a patient’s own iPSC-derived neurons. Fourth, we used nonviral particles to deliver transient Cas9 RNP or mRNA, rather than viral vectors delivering genetically encoded Cas9. This avoided indefinite Cas9 expression that would have obscured the prolonged editing time course, and avoided exogenous DNA episomes that could be aberrantly integrated into long-lived DSBs. Nonviral delivery to postmitotic human neurons has long been a challenge for the field; utilizing recent advances in VLP and LNP technology to overcome this barrier was crucial to studying neuronal DNA repair accurately.

Our approach also had several limitations, which could be addressed with future follow-up studies. First, since the DSB detection assays available to us were endpoint assays, we could not definitively test whether multiple cycles of cutting and resealing occur within the same cell. Second, while certain siRNAs increased indel efficiency compared to NT siRNA, overall indel efficiency with LNPs was still fairly low and more variable batch-to-batch. We used this proof-of-concept siRNA platform mainly as a genetic tool to investigate our hypotheses about DNA repair factors in neurons. Any groups aiming to advance these all-in-one LNPs as a therapeutic tool should optimize the lipid formulation, the species and ratios of siRNAs, and the timing of knockdown relative to editing. Third, while our studies were conducted in postmitotic human neurons, it is unknown how our findings will translate to aged/diseased neurons in patients, or nonhuman neurons in animal models. Future studies could investigate the timing and repair of edits in rodent/primate neurons, and potentially in *ex vivo* primary human neurons. Finally, it is very likely that other untested sgRNAs, and/or other nucleases, will have different DSB repair dependencies than the ones revealed by this study. In follow-up studies, we plan to pair our platform with higher-throughput methods such as CRISPR interference to find optimal repair modifications for particular sgRNAs of interest, and study neuronal responses to other genome editing enzymes.

In summary, examining how postmitotic neurons respond to CRISPR perturbations uncovered new considerations for safety and efficacy, and new avenues for controlling CRISPR repair outcomes. The genome modification toolkit contains several tools to perturb DNA, but we are just beginning to develop tools that ensure proper repair. Those tools will be crucial for unlocking the full potential of therapeutic genome editing.

## EXTENDED DATA

**Extended Data Figure S1:**
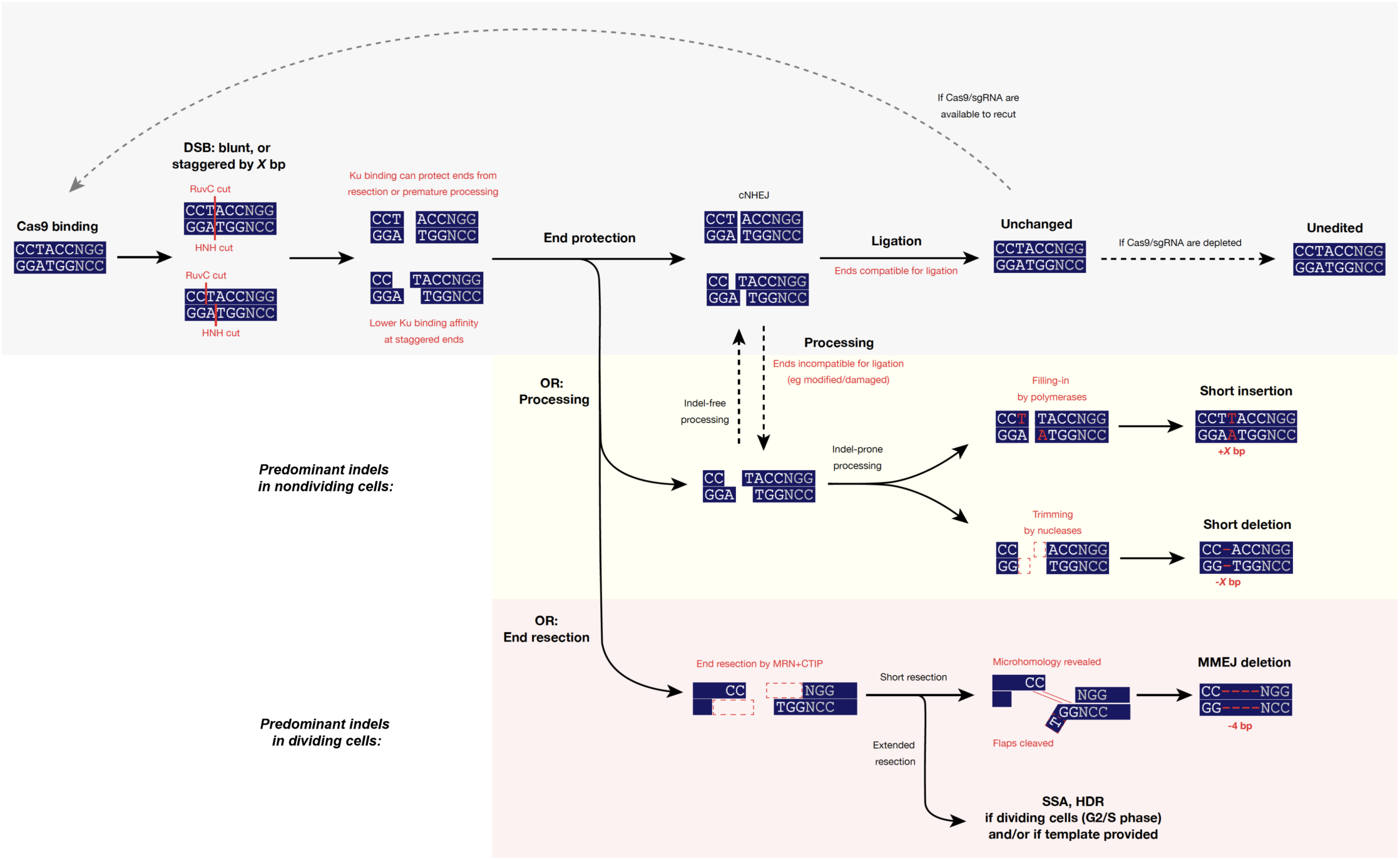
Schematic of how DSB repair pathways determine the CRISPR editing outcome. Cas9 induces a blunt or staggered DSB, depending on where the RuvC domain cleaves (Shou et al, Mol Cell, 2018. PMID: 30033371). The exposed DNA ends are then subjected to either end protection or end resection (or other processing). End protection generally leads to cNHEJ. If the protected ends are still chemically compatible for ligation, cNHEJ often ligates them fidelitously, yielding an unchanged sequence which can be re-cut by any remaining Cas9 RNP. If the protected ends are not compatible for ligation, or if end protection was outcompeted by processing machinery such as polymerases and nucleases, then NHEJ processing can occur (Stinson et al, Mol Cell, 2020. PMID: 31862156). This processing sometimes introduces indels. In dividing cells, end resection often outcompetes end protection, leading to resection-dependent pathways such as MMEJ, HDR and SSA. Resection-dependent pathways can cause indels (MMEJ/SSA) or templated repair (HDR).

**Extended Data Figure S2:**
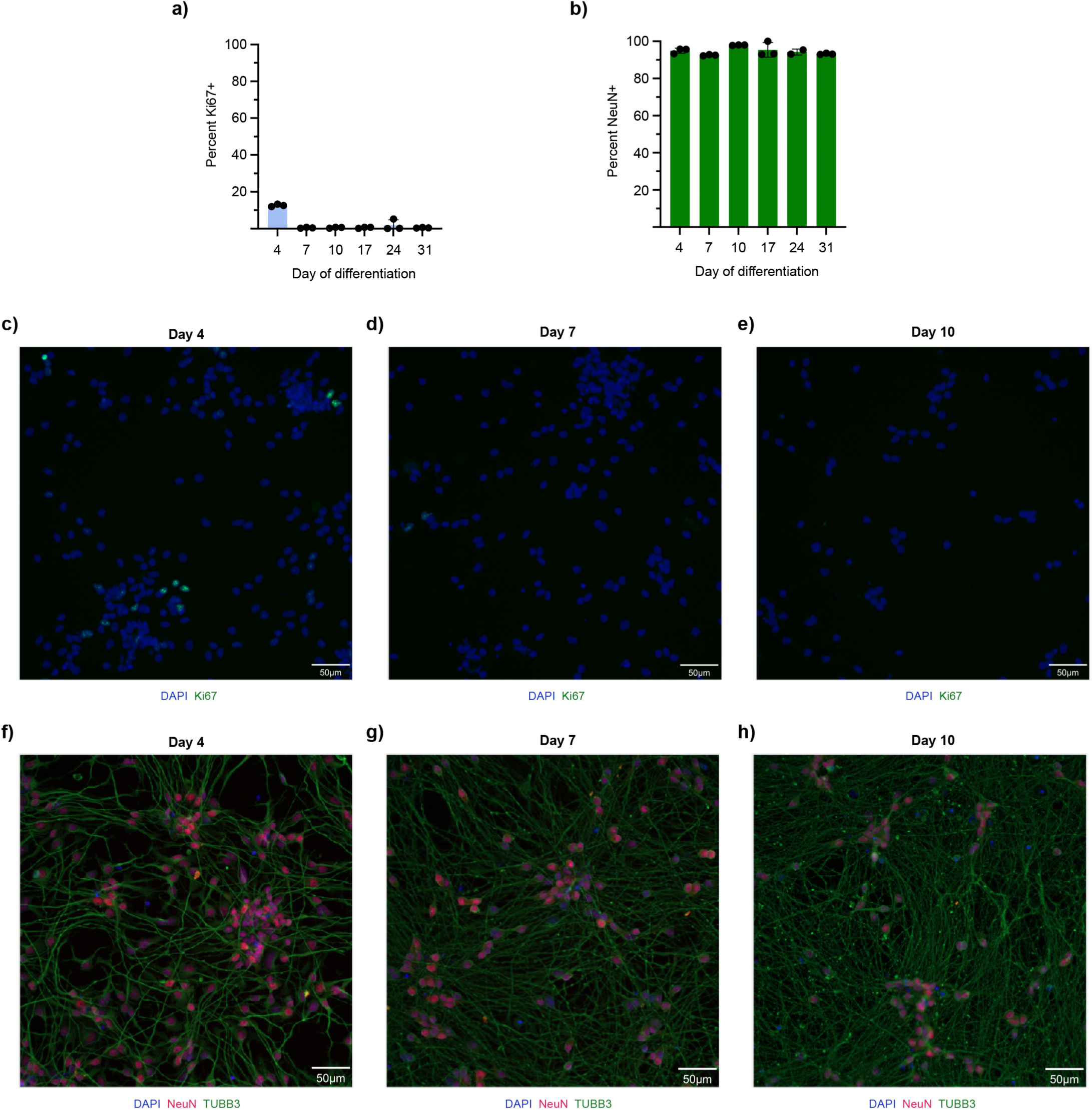
Characterizing the purity of the neuronal differentiation. **a)** By Day 7 of differentiation, less than 1% of cells are proliferative (Ki67+). Bars show what percentage of DAPI+ nuclei were Ki67+, averaged across 3 replicate wells. HCS Studio SpotDetector. **b)** By Day 4 of differentiation, 95% of cells express a neuron-specific marker (NeuN+). Bars show what percentage of DAPI+ nuclei were NeuN+, averaged across 3 replicate wells. CellProfiler. For a-b: Each dot is one replicate well, totaled across 13 non-overlapping fields per well. Error bars show SEM. **c-e)** Representative ICC images showing DAPI and Ki67 staining from Days 4/7/10 of differentiation; quantified in a. **f-h)** Representative ICC images showing DAPI, NeuN, and TUBB3 staining from Days 4/7/10 of differentiation; quantified in b. TUBB3 is another marker of mature neurons. **Note for c-h**: HCS Studio software baked the scale bar annotations into the output montages, and only the “merged” panel of each montage is shown here. Differences in font size and bar thickness are simply due to differences in dimensions between two-panel (DAPI+Ki67) and three-panel (DAPI+NeuN+TUBB3) montages. Scale bar lengths remain accurate for each panel.

**Extended Data Figure S3:**
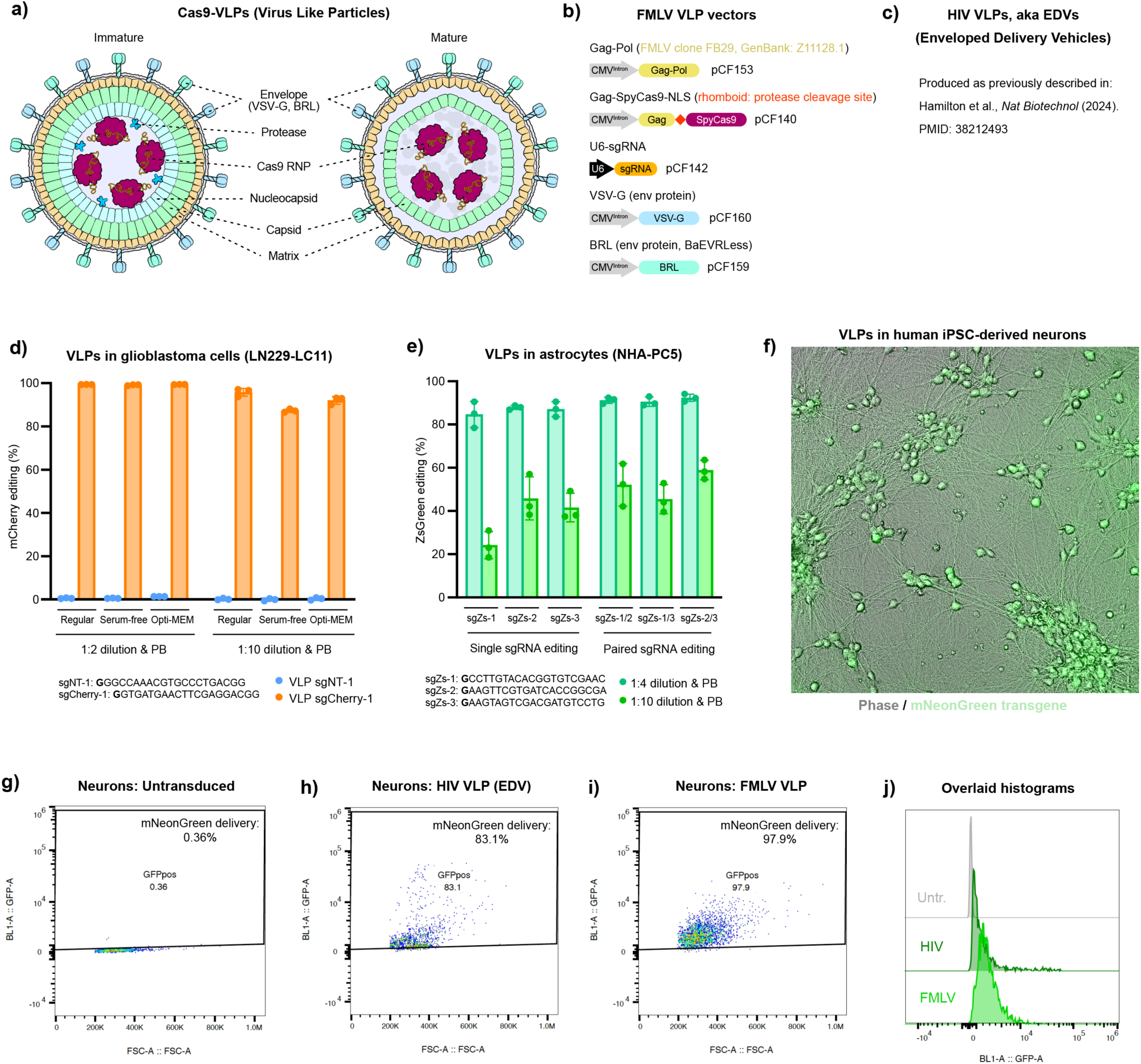
Establishing VLP delivery of Cas9 to human postmitotic neurons. **a)** Schematic depicting the components of virus-like particles. Matrix, capsid, and nucleocapsid are part of the Gag polyprotein. VSV-G and BRL are envelope (env) proteins for pseudotyping to mediate broad and efficient cellular transduction. **b)** Maps and nomenclature of optimized FMLV VLP vectors. **c)** Vectors used to produce HIV VLPs, also known as enveloped delivery vehicles (EDVs), were previously described in Hamilton et al., Nat Biotechnol, 2024. PMID: 38212493. **d)** Assessment of editing efficiency with optimized FMLV VLPs in glioblastoma cells. Monoclonal mCherry-expressing glioblastoma cells (LN229-LC11) were transduced with the indicated VLPs harvested in regular growth media, serum-free growth media, or Opti-MEM. Target cells were transduced at the indicated VLP dilution, with addition of polybrene (PB, 5 μg/ml). At day six post-transduction, mCherry editing efficiency (mCherry-) was assessed by flow cytometry. Non-transduced cells were used for normalization. VLP sgCherry-1: CRISPR-Cas9 VLP containing a previously validated mCherry-targeting sgRNA (Knott et al, eLife, 2019. PMID: 31397669). VLP sgNT-1: CRISPR-Cas9 VLP containing a non-targeting control sgRNA. Error bars indicate standard deviation. **e)** Assessment of editing efficiency with optimized FMLV VLPs in astrocytes. Normal human astrocytes expressing ZsGreen (NHA-PC5), and previously treated with puromycin-targeting VLPs (Tan et al, Cell Reports, 2023. PMID: 37917583), were transduced with sgZsGreen-targeting VLPs (harvested in regular growth media) at the indicated dilution, with addition of polybrene (PB). Cells were either transduced with a single VLP to generate indels or with a mixture of two VLPs to induce a deletion in ZsGreen. At day six post-transduction, ZsGreen editing efficiency (ZsGreen-) was assessed by flow cytometry. Non-transduced cells were used for normalization. VLP sgZs-1/2/3: CRISPR-Cas9 VLPs containing ZsGreen-targeting sgRNAs. Error bars indicate standard deviation. **f)** Optimized FMLV VLPs efficiently transduced human iPSC-derived neurons. Representative microscopy image of neurons after transduction with Cas9 VLPs co-encapsulating an mNeonGreen transgene. **g-j)** Our optimized FMLV VLPs and HIV VLPs (EDVs) both transduced human iPSC-derived neurons efficiently. Flow cytometry 1-week post-transduction with no VLP (untransduced), HIV VLPs, or optimized FMLV VLPs shows up to 97% transduction efficiency. Dosage: 2 µL VLP per 100 µL media.

**Extended Data Figure S4:**
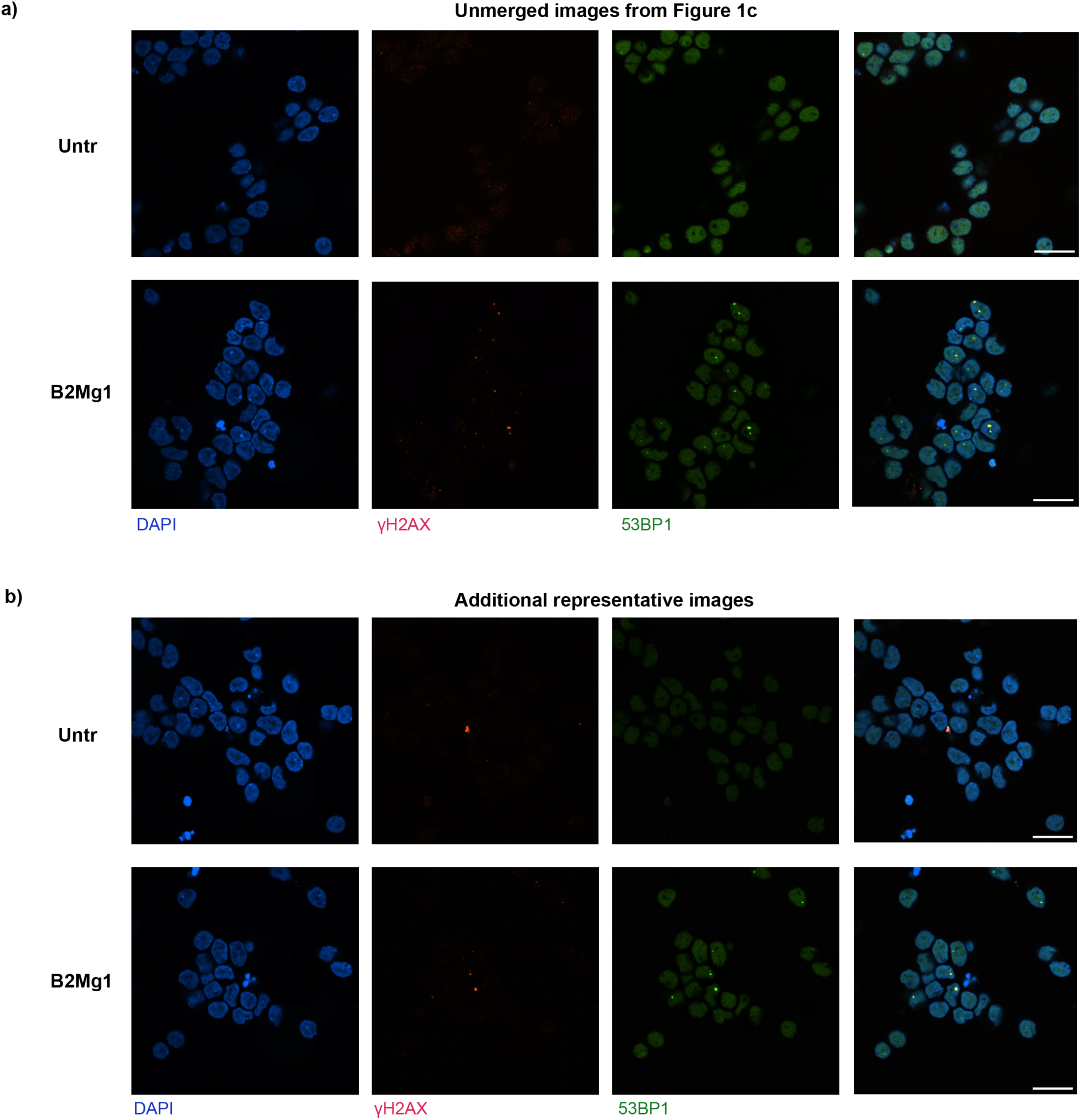
Cas9-VLPs induce DSBs in human postmitotic neurons. **a)** Unmerged panels from Figure 1c, showing DSBs induced by Cas9-VLPs in human iPSC-derived neurons, compared to age-matched untransduced neurons. For a-b: Neurons transduced 2 weeks into differentiation, and imaged 3 days post-transduction. DSBs are co-labeled by markers γH2AX (red) and 53BP1 (green). Dose: 1 µL FMLV VLP per 100 µL media. Scale bar is 20 µm. **b)** Additional representative ICC images showing DSBs induced by Cas9-VLPs in human iPSC-derived neurons, compared to age-matched untransduced neurons.

**Extended Data Figure S5:**
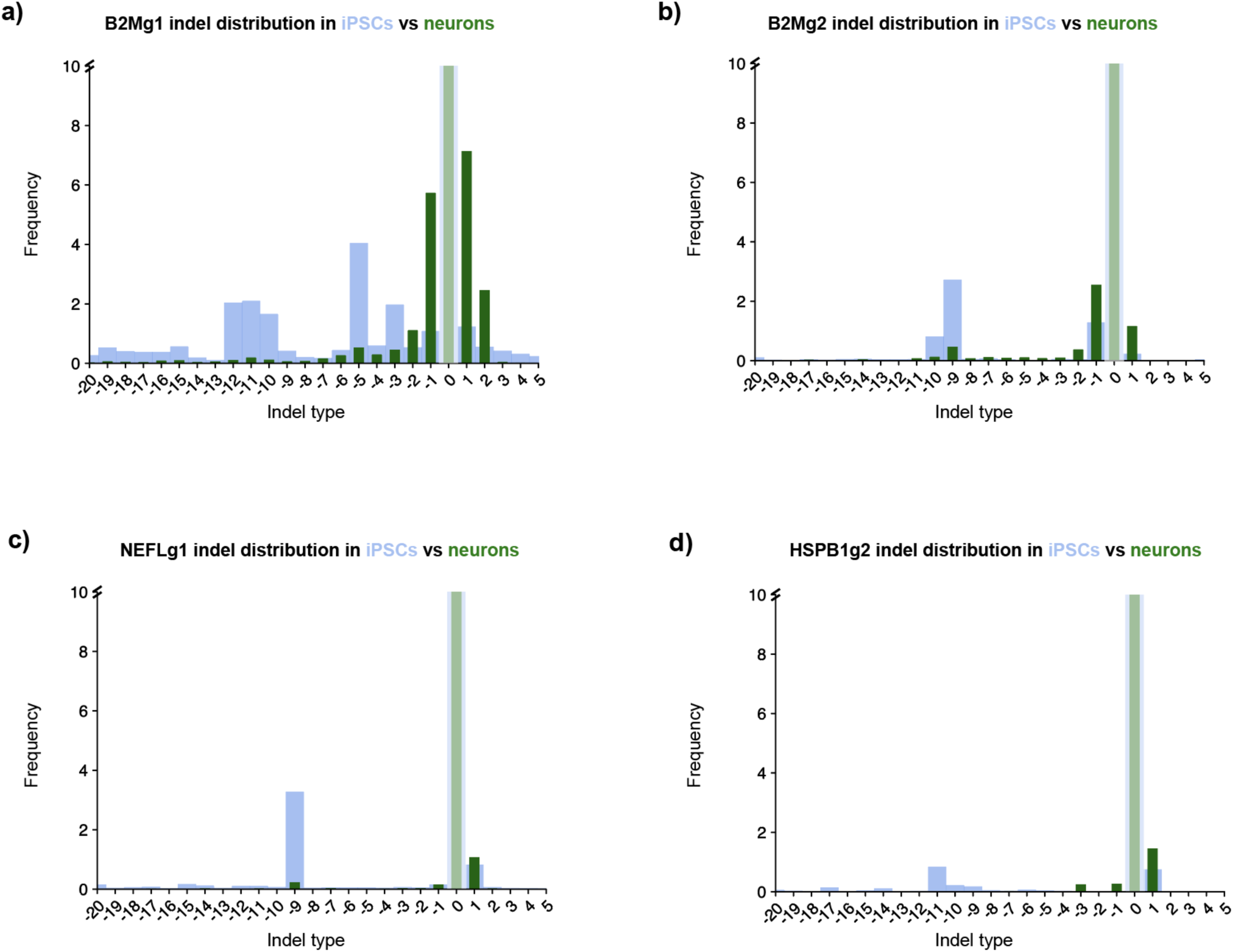
CRISPR editing outcomes differ in postmitotic neurons compared to isogenic dividing cells. **a-d)** For each of four separate sgRNAs, CRISPR editing outcomes differ between nondividing neurons and dividing iPSCs. Despite differences in which indel outcomes each sgRNA was amenable to overall, in each case, the MMEJ-like deletions were predominant in iPSCs whereas the NHEJ-like small indels were predominant in neurons. Dose: 2 µL VLP (FMLV) per 100 µL media. Average of 6 replicate wells, transduced in parallel. Genomic DNA was harvested 5 days post-transduction, processed for amplicon-NGS, then analyzed by CRISPResso2.

**Extended Data Figure S6:**
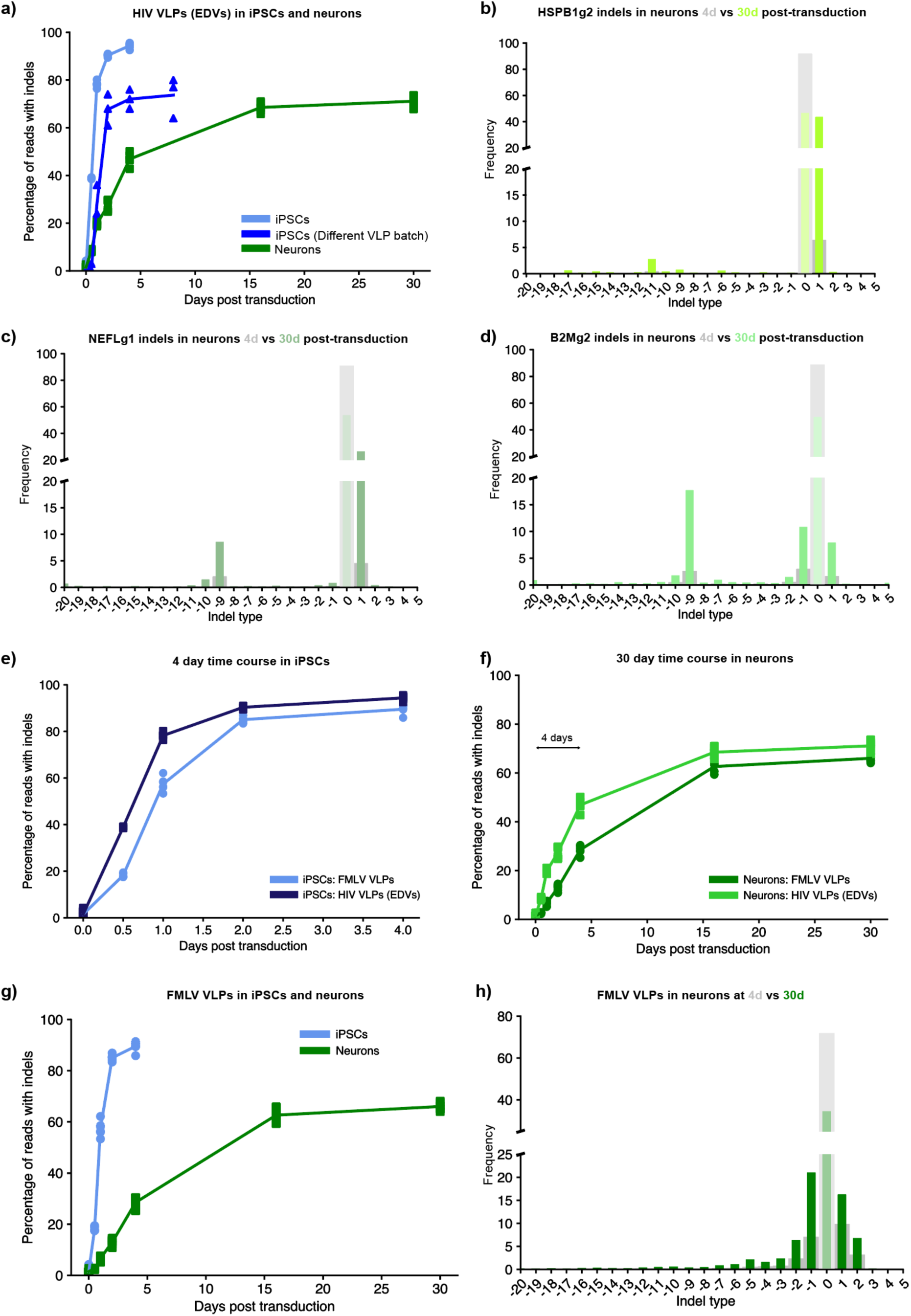
Cas9-VLP-induced indels accumulate for weeks post-transduction in neurons. **a)** Even in experiments where iPSCs plateaued at a lower editing efficiency, they still reached that plateau sooner than neurons. Regraphed data from Figure 2a, and overlayed data from a separate experiment with a less-efficient batch of VLPs. In this experiment, iPSCs plateaued at 60-70% indels instead of 90%+, but still reached that plateau within ∼4 days. **b-d)** For three additional sgRNAs, despite differences in which indel outcomes each sgRNA was amenable to overall, all available indel outcomes at the 4 day timepoint increased in prevalence by the 30 day timepoint. Dose: 1 µL VLP (FMLV) per 100 µL media. **e-h)** The time course of indel accumulation was reproduced very comparably between FMLV and HIV based Cas9-VLPs. With either delivery particle, indels plateaued within 4 days post-transduction in iPSCs (e), but continued to increase for up to 16 days post-transduction in neurons (f). The overlaid time courses look very similar with FMLV VLPs (g-h) compared to HIV VLPs (Figure 2a, 2c). Dose: 2 µL VLP per 100 µL media.

**Extended Data Figure S7:**
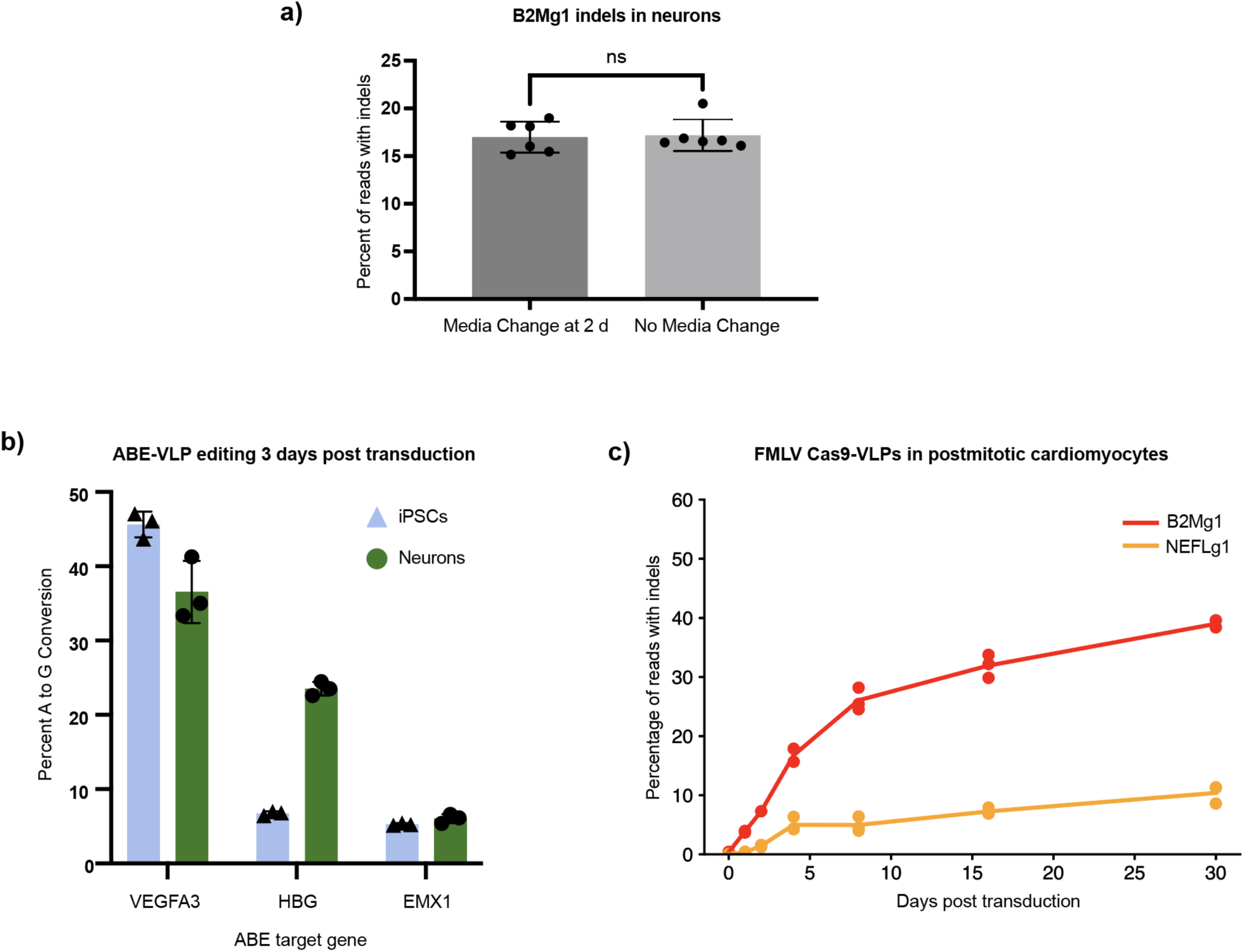
Testing additional hypotheses about the prolonged editing time course in neurons. **a)** The prolonged indel accumulation in neurons is **not** driven by residual VLP in the media. Replacing the media after 2 days post-transduction (as is required for iPSCs) did not significantly affect neuron editing efficiency at 4 days post-transduction. Notable because the steepest increase in indels in neurons typically occurs over the first 4 days post-transduction. Unpaired t test, ns = not significant (p>0.05). 6 replicate wells per condition, transduced in parallel. Dose: 1 µL VLP (FMLV) per 100 µL media. **b)** ABE-VLPs confirm that the slow indel accumulation we observed is **not** a product of deficient VLP delivery to neurons. When the same HIV VLPs are used to deliver adenine base editors (ABEs) instead of Cas9, neurons can match and even exceed the editing efficiency of iPSCs, within only 3 days post-transduction. Error bars show SEM; 3 replicate wells per condition, transduced in parallel. Dose: 4 µL VLP (HIV) per 100 µL media – but these VLPs were half as concentrated as normal, since VLPs were harvested from 3 10 cm plates per batch instead of 6 but still resuspended to the same volume. Therefore, equivalent to a 2 µL VLP dose from normal batches. ABE-VLP cloning protocol is described in the fourth tab of Supplemental Table 3. **c)** Postmitotic iPSC-derived cardiomyocytes (CMs) also show a weeks-long accumulation of indels, for two different sgRNAs. At day 30+ of differentiation, after lactate purification to select for postmitotic CMs, CMs were transduced with 1 µL FMLV VLP per 100 µL media. 3 replicate wells per condition, transduced in parallel. CRISPResso2 analysis of amplicon-NGS. CMs were generated from WTC background iPSCs using the protocol described in Lian et al, Nat Protoc, 2013 (PMID: 23257984).

**Extended Data Figure S8:**
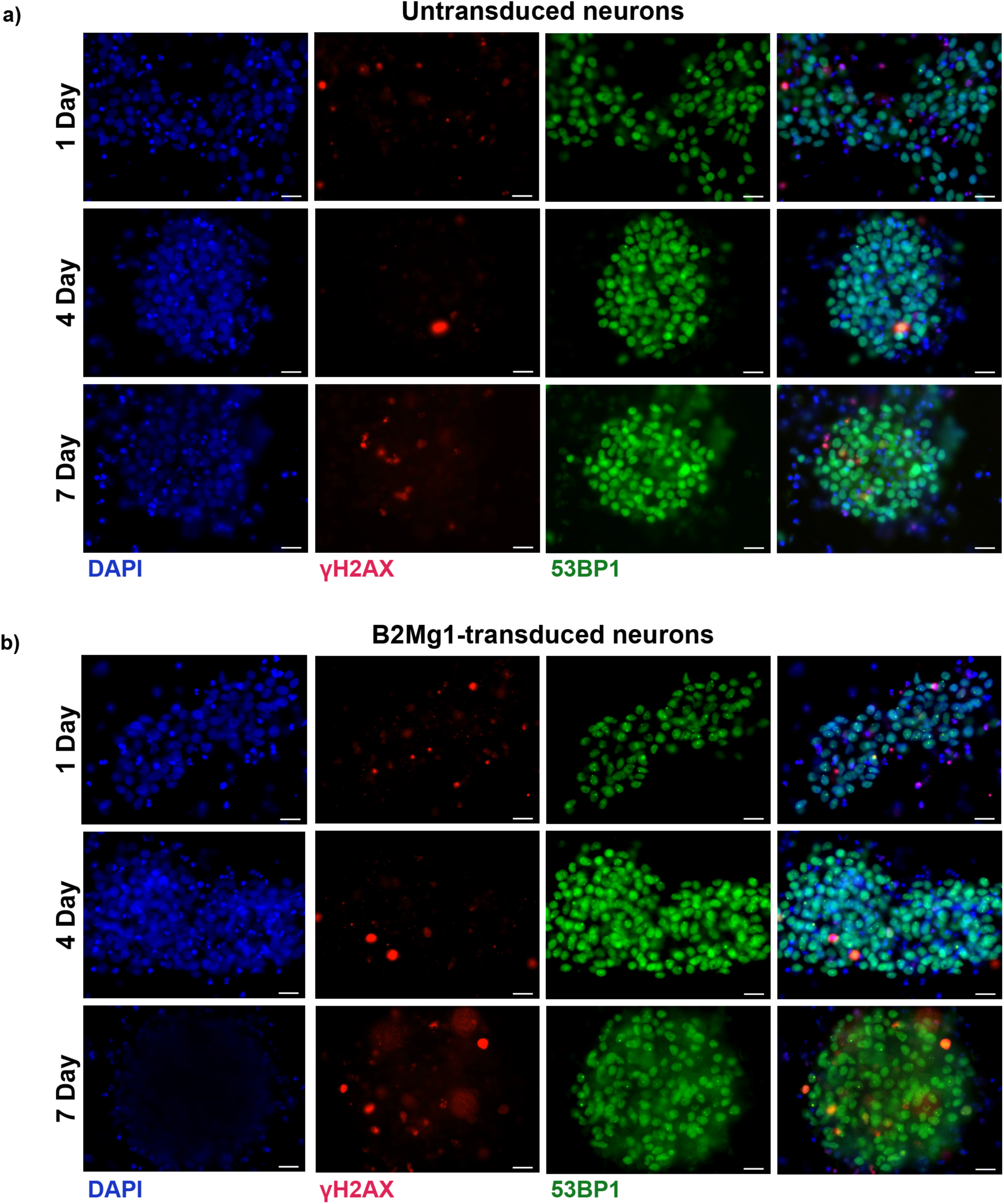
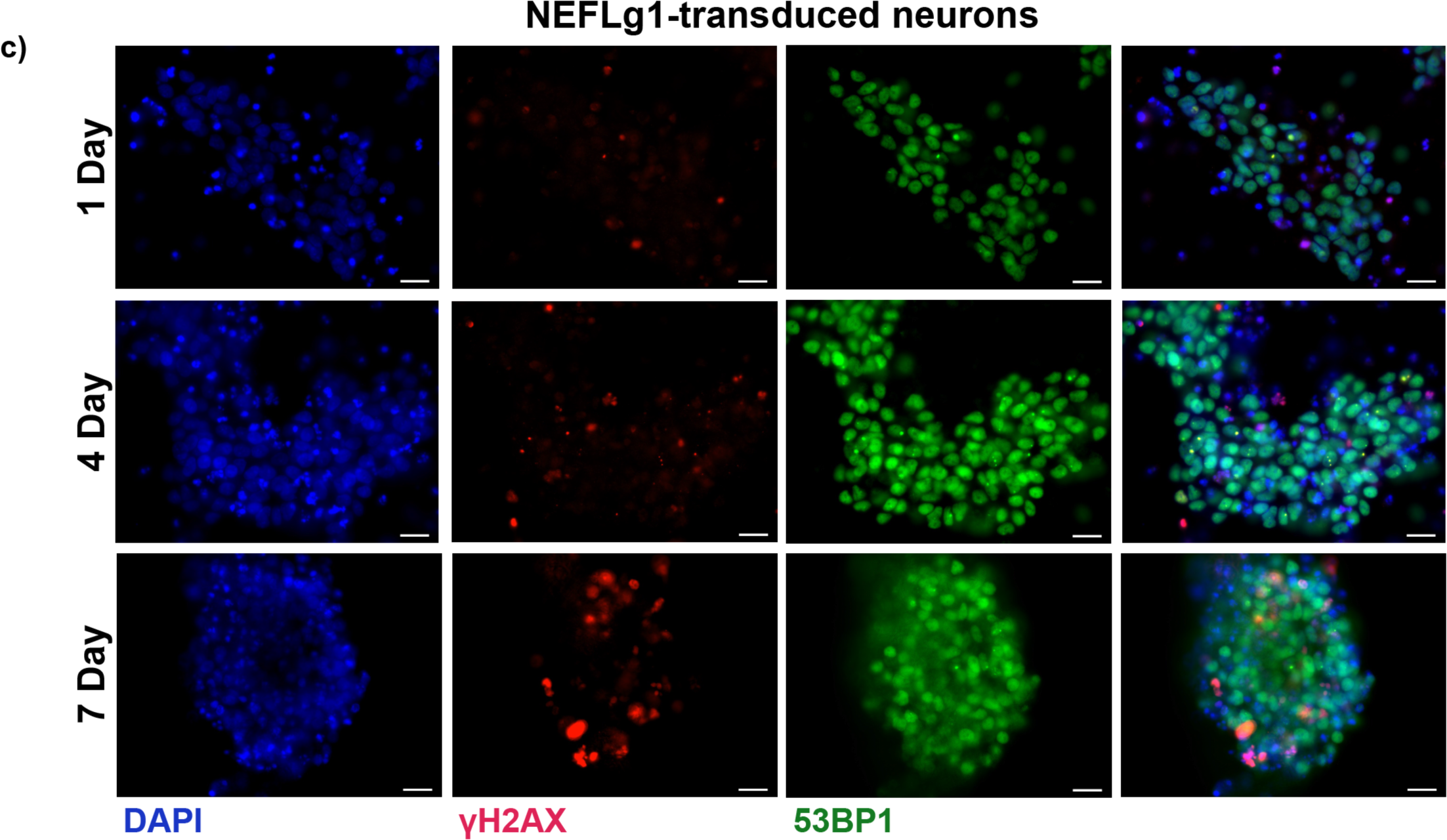
Cas9-induced DSB repair signals persist in neurons for at least one week post-transduction. **a-c)** DSB repair markers over time in untransduced (a), B2Mg1-transduced (b), and NEFLg1-transduced (c) neurons. DSBs are co-labeled by ICC markers γH2AX (red) and 53BP1 (green). Dose: 1 µL FMLV VLP per 100 µL media. Neurons were fixed at 1,4, or 7 days post-transduction as labeled. One representative image from each condition is shown. Transduction was 2 weeks into differentiation. Scale bar is 20 µm. Same experiment as Figure 2d-e, but now showing unmerged panels individually, and including additional conditions (timepoints, sgRNAs). Therefore, the merged panels for untransduced and B2Mg1-transduced at 1 day and 7 days are the same as in Figure 2d-e, but uncropped here.

**Extended Data Figure S9:**
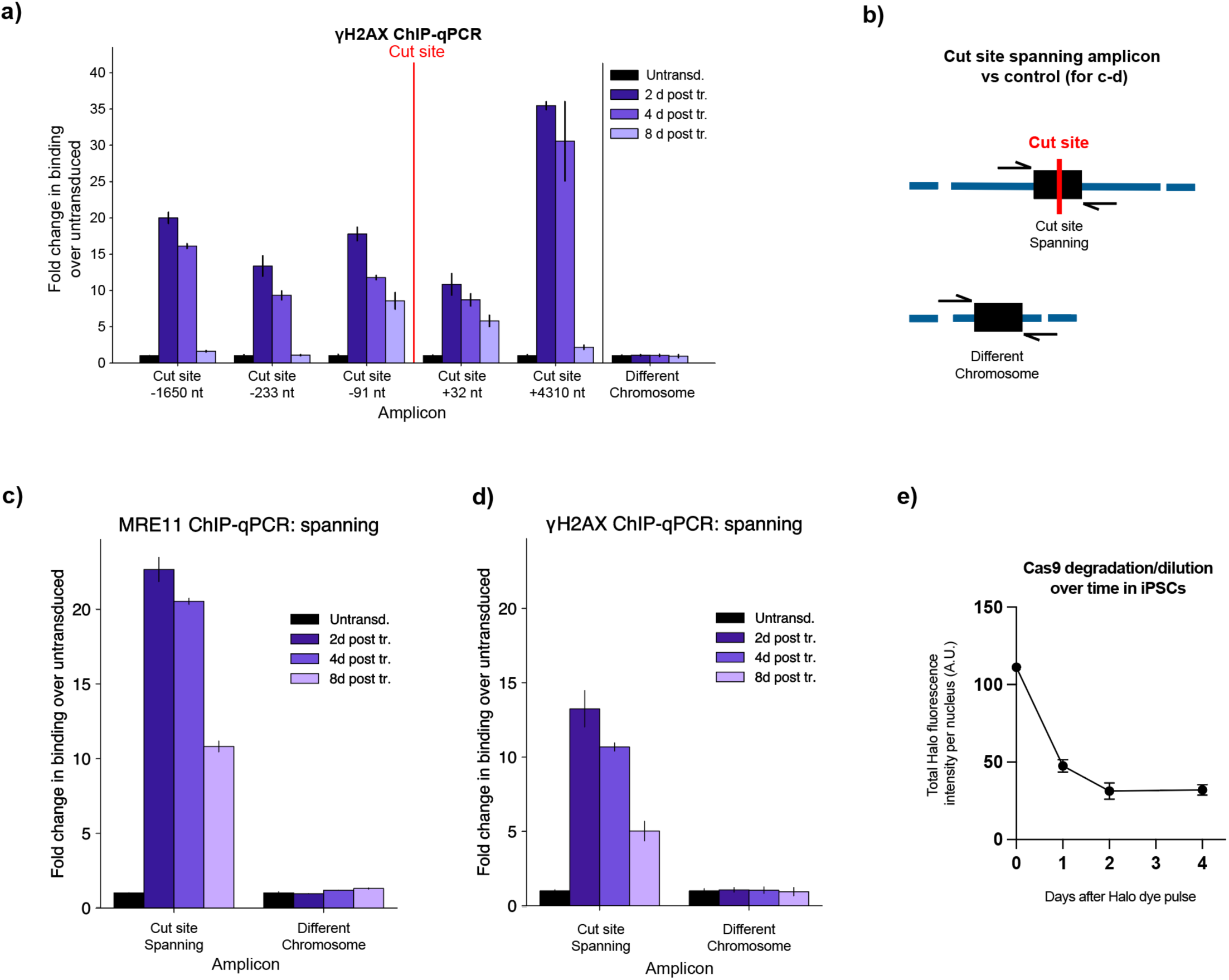
DSB repair signals remain detectable at the cut site for at least 8 days post-transduction. **a)** ChIP-qPCR for γH2AX binding at various distances from the cut site over time. Same conditions as Figure 2f, but with γH2AX antibody instead of Mre11. **b)** Schematic illustrating our strategy to detect cut-and-resealed loci by using a ChIP-qPCR amplicon that spans across the cut site. Repair protein binding suggests that the locus had been cut, and successful PCR amplification suggests that the cut has since been resealed. Note: however, it remains ambiguous whether these loci were sealed with or without an indel. **c-d)** Some loci have been resealed as early as 2 days post-transduction. ChIP-qPCR using the spanning amplicon to detect cut-and-resealed loci, with both Mre11 (c) and gH2AX (d). Same procedures as Figure 2f and S9a, but using different amplicons (cut site spanning, and different chromosome control). **e)** Cas9 protein in iPSCs gets quickly diluted and/or degraded to background levels within 2 days post-transduction; therefore, these neuron ChIP-qPCR data cannot be compared to iPSCs. Pulse-chase to track degradation of Halo-tagged Cas9 in iPSCs. First, iPSCs (with/without lentivirally integrated Halo-Cas9 and B2Mg1) were seeded onto glass-bottom 96-well plates with ∼2,000 cells per well. iPSCs were pulsed with 40 µM fluorescent Halo ligand (Promega HaloTag-JF549, cat. #GA1110) for 1 hour, then washed with fresh media 3 times to prevent newly translated Cas9 protein from being labeled. iPSCs were then chased with 2 µM of an unlabeled Halo ligand (Promega ent-HaloPROTAC3, cat. #GA4110) as a binding competitor. Nuclei were labeled with NucBlue (ThermoFisher, cat. #R37605) 20 min before live cell imaging on the Image Xpress Confocal Microscope. Halo fluorescence signal was measured at several timepoints to track the degradation/dilution of the pulse-labeled Cas9 molecules over time. Analyzed in CellProfiler. 8 replicate wells; error bars show standard deviation.

**Extended Data Figure S10:**
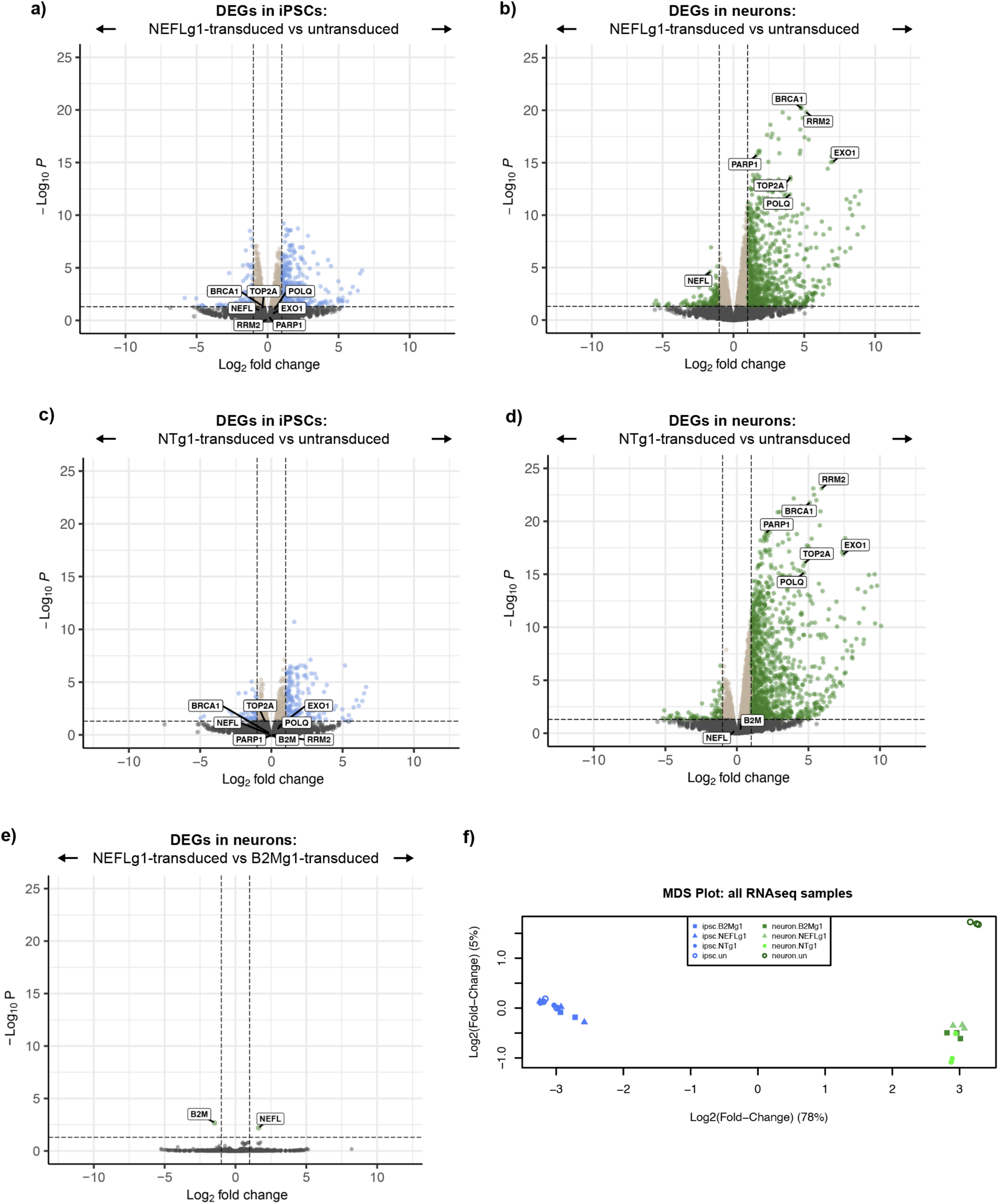
Neuronal transcriptional response to Cas9-VLP is very consistent across three different sgRNAs. **a-d)** The neuron-specific transcriptional response to Cas9-VLPs was replicated by two additional sgRNAs, NEFLg1 (a-b) and NTg1 (c-d), besides B2Mg1 shown in Figure 3a-b. Neurons have more DEGs overall upon transduction, and the most significant of these DEGs are enriched for DNA repair genes. Same parameters as Figure 3a-b, but with different sgRNAs. Note: *NEFL* is not expressed in iPSCs, so its expression is not expected to decrease upon NEFLg1 editing in iPSCs. **e)** The only two DEGs between B2Mg1-edited and NEFLg1-edited neurons are *B2M* and *NEFL* respectively. This reinforces the consistency of the neuronal transcriptional response across different sgRNAs. **f)** The transcriptional profile of NTg1-treated neurons is more similar to B2Mg1-edited and NEFLg1-edited neurons than to untransduced neurons. Therefore, at least some component of the neuronal response to Cas9-VLPs may be DSB-independent. Multidimensional scaling (MDS) plot visualizing similarity between the various RNAseq samples. As indicated in this plot, all RNA-seq analysis in this study was conducted on 3 replicate samples per condition, transduced in parallel; 24 total RNA-seq samples.

**Extended Data Figure S11:**
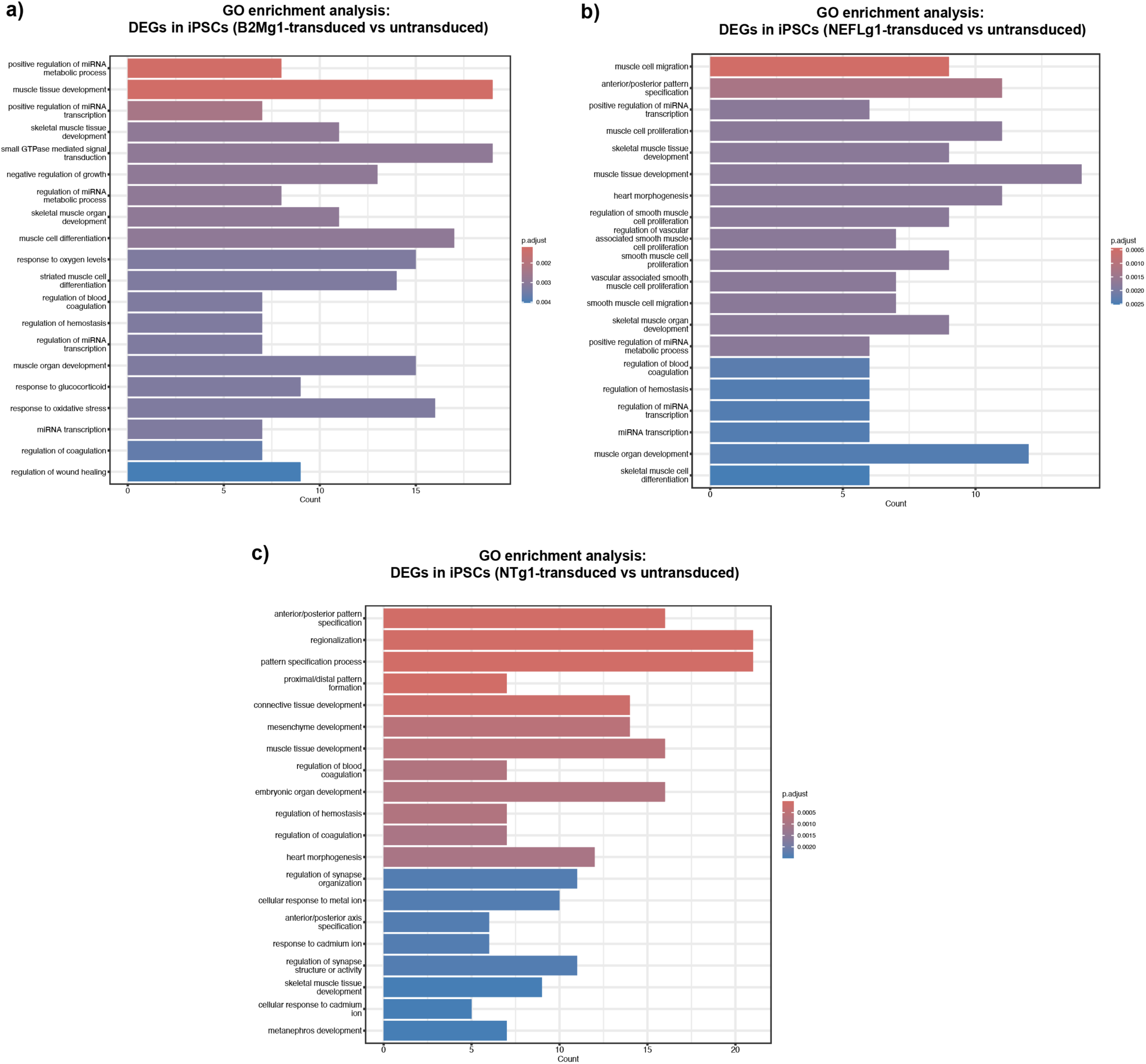
DNA repair genes are not enriched in the DEGs of transduced iPSCs. **a-c)** Gene ontology (GO) analysis shows no enrichment for DNA repair genes in the DEGs of B2Mg1-transduced (a), NEFLg1-transduced (b), or NTg1-transduced (c) iPSCs, relative to untransduced iPSCs. Showing the top 20 GO terms in each comparison. Bar length indicates number of DEGs that fall into each GO category. Color indicates significance of adjusted p-value.

**Extended Data Figure S12:**
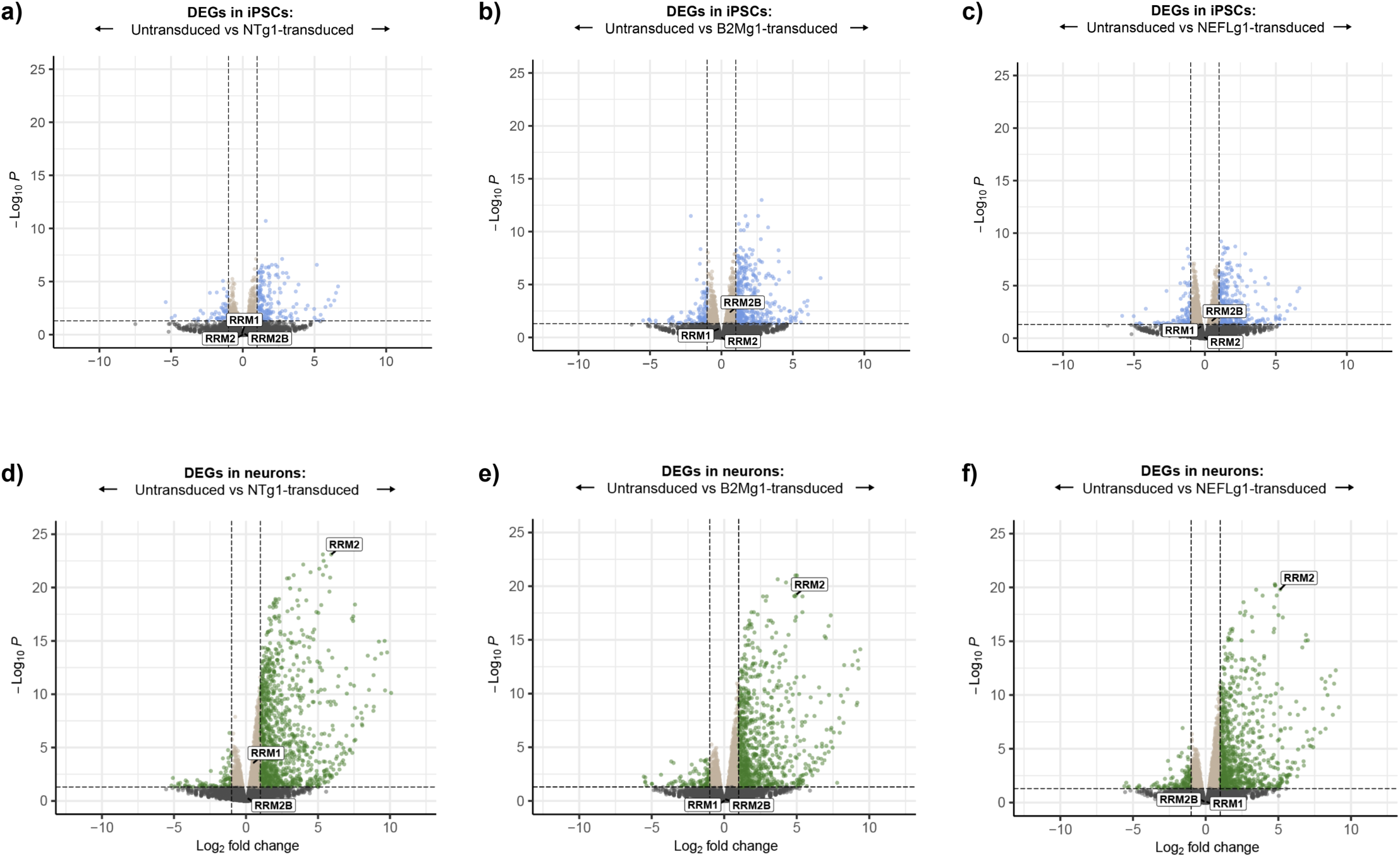
Transcriptional response of RNR subunits in neurons compared to iPSCs. **a-c)** In iPSCs, non-targeting Cas9 (a) does not affect transcription of any RNR subunits. However, both of the cutting Cas9-VLPs (b-c) significantly induce transcription of *RRM2B*, the canonically DSB-responsive subunit of RNR. The other two subunits are unaffected. **d-f)** In neurons, the canonically S-phase-restricted *RRM2* is one of the most significantly upregulated genes in all 3 transduced conditions, including with non-targeting Cas9.

**Extended Data Figure S13:**
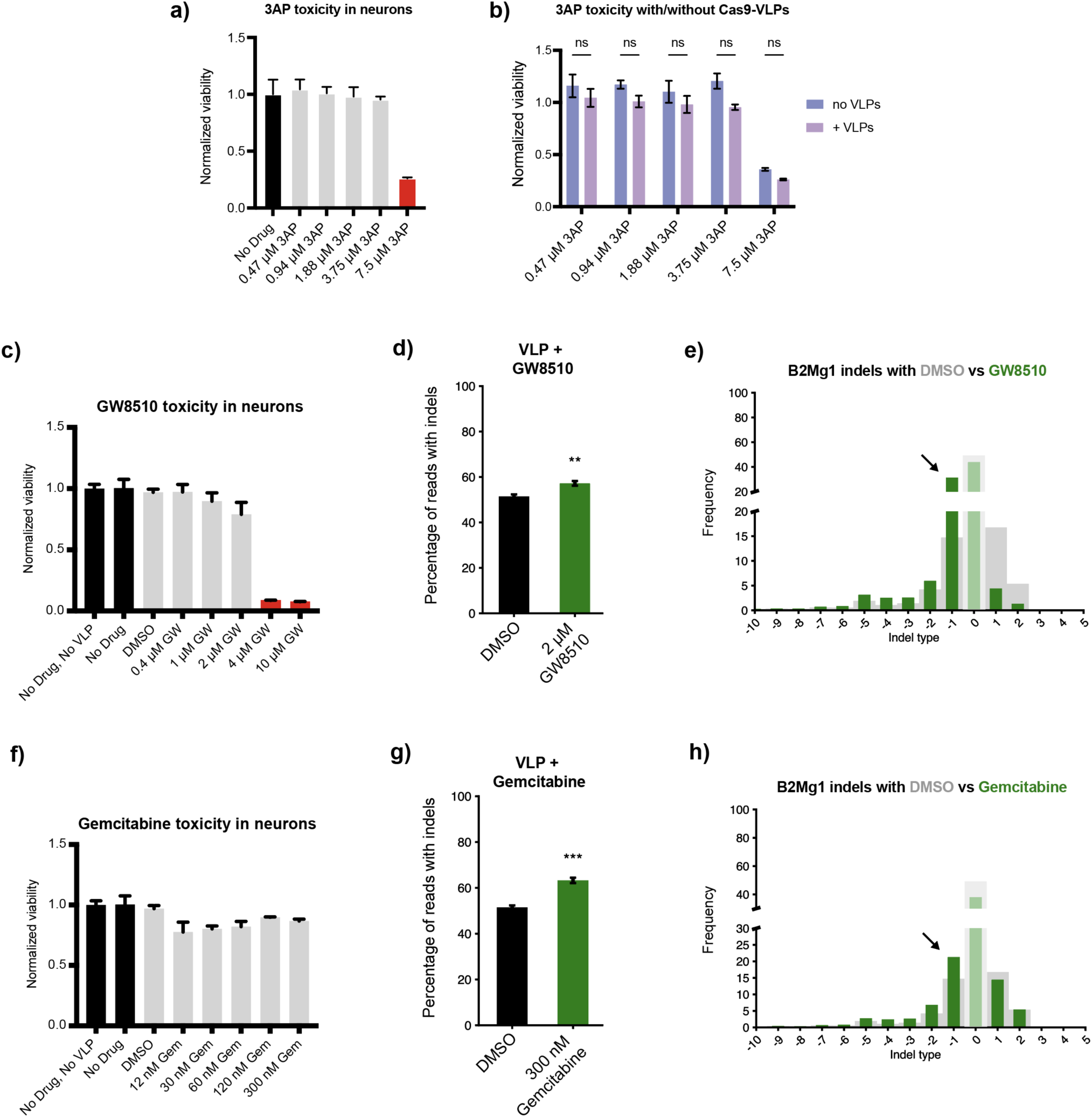
Inhibiting RNR changes editing outcomes in neurons. **a)** Toxicity of escalating doses of RRM2 inhibitor 3AP in neurons. Maximum tolerable dose was 3.75 µM. For a/c/f: Tolerability threshold was arbitrarily set to 0.75 or above, corresponding to less than a 25% reduction in viability. PrestoBlue viability assay at 8 days post-transduction, normalized to age-matched untreated neurons on the same plate. 3 replicate wells per condition, treated in parallel; error bars show SEM. **b)** Toxicity of escalating 3AP doses in neurons with or without Cas9-VLPs. Optimal 3.75 µM dose remains nontoxic even with Cas9-VLPs (1 µL FMLV) inducing DNA damage. Error bars show SEM. Two-factor ANOVA; ns = not significant (p>0.05). **c)** Toxicity of escalating doses of RRM2 inhibitor GW8510 in neurons, alongside Cas9-VLP treatment. Maximum tolerable dose was 2 µM. **d-e)** GW8510 cotreatment of B2Mg1-edited neurons increases indels overall (d), boosting deletions specifically, and roughly doubles the frequency of single-base deletions (e). Replicated the effects of RRM2 inhibitor 3AP from Figure 3g-h. Dose: 1 µL FMLV VLP per 100 µL media, and maximum tolerable dose of GW8510. Indels measured 8 days post-transduction. For d, error bars show SEM. One-Factor ANOVA, ** p<0.005. For d-e, 6 replicate wells per condition treated in parallel. **f)** Toxicity of escalating doses of RRM1 inhibitor gemcitabine in neurons, alongside Cas9-VLP treatment. Maximum tested dose was 300 nM, and still tolerable. **g-h)** Gemcitabine co-treatment of B2Mg1-edited neurons increases indels overall (g), boosting deletions specifically (h). Replicated the effect of RRM2 inhibitor 3AP from Figure 3g-h. Dose: 1 µL FMLV VLP per 100 µL media, and maximum tolerable dose of gemcitabine. Indels measured 8 days post-transduction. For g, error bars show SEM. One-Factor ANOVA, *** p<0.0005. For g-h, 6 replicate wells per condition treated in parallel.

**Extended Data Figure S14:**
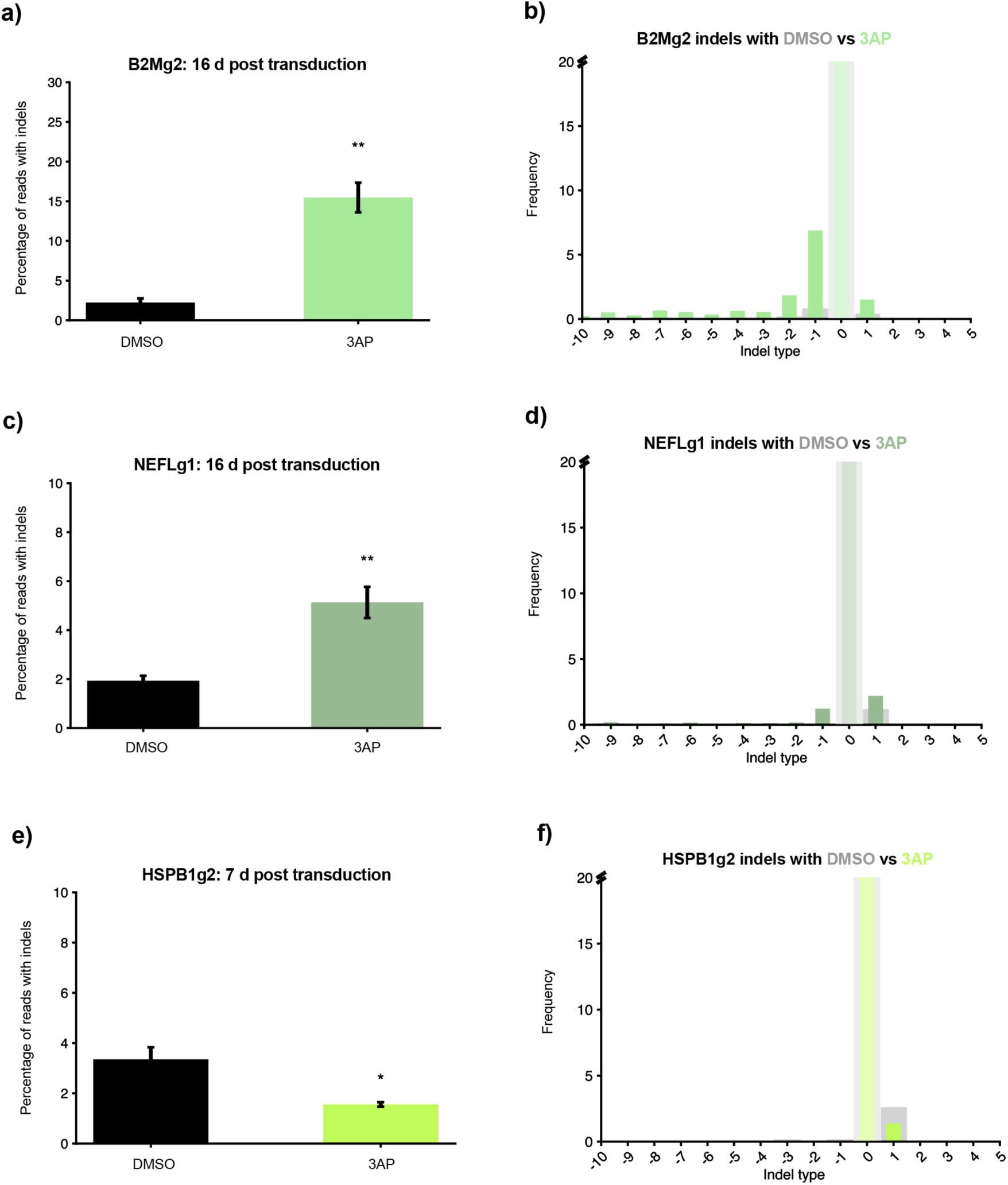
RNR inhibition affects neuron editing outcomes in an sgRNA-dependent manner. **a-d)** For B2Mg2 (a-b) and NEFLg1 (c-d), 3AP treatment significantly increases indels, but without the same selectivity for deletions as seen for B2Mg1. 6 replicate wells per condition, treated in parallel. **e-f)** For HSPB1g2, 3AP treatment significantly decreases indels instead of increasing them. This is consistent with its intrinsic indel distribution, which appears relatively impermissible to deletions. 6 replicate wells per condition, treated in parallel. For a/c/e: One-Factor ANOVA, ** p<0.005, * p<0.05. For a-f: 1 µL dose, FMLV VLPs.

**Extended Data Figure S15:**
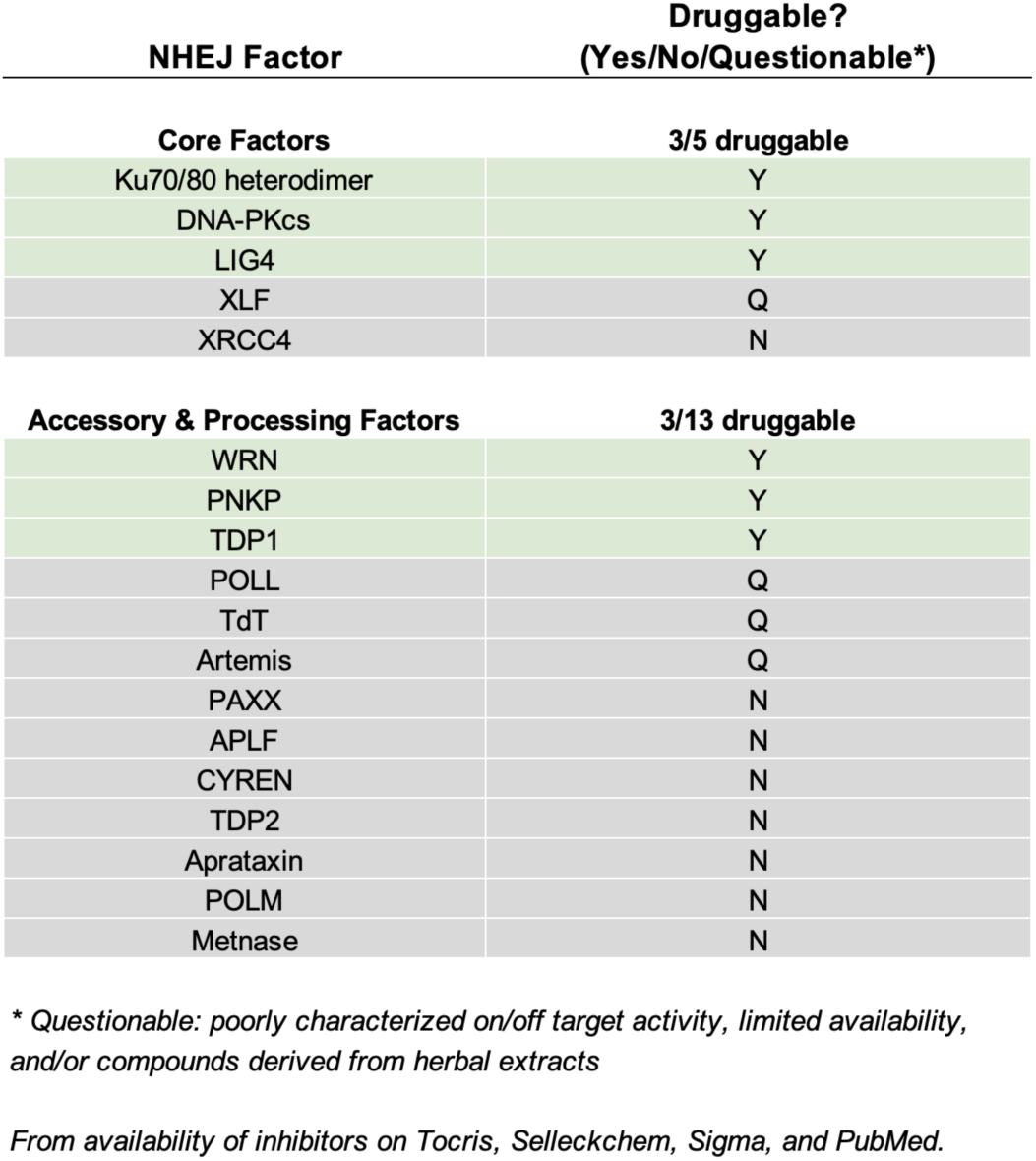
NHEJ pathway exemplifies that many DNA repair factors are not reliably targetable with small molecule inhibitors. Two-thirds of the factors in the NHEJ pathway (Stinson et al, Annu Rev Biochem, 2021. PMID: 33556282) are not reliably druggable by small molecule inhibitors. Determined by searching for availability of inhibitors on Tocris, Selleckchem, Sigma, and PubMed, as of 2023. This simply demonstrates how many DNA repair factors are not readily druggable at the protein level.

**Extended Data Figure S16:**
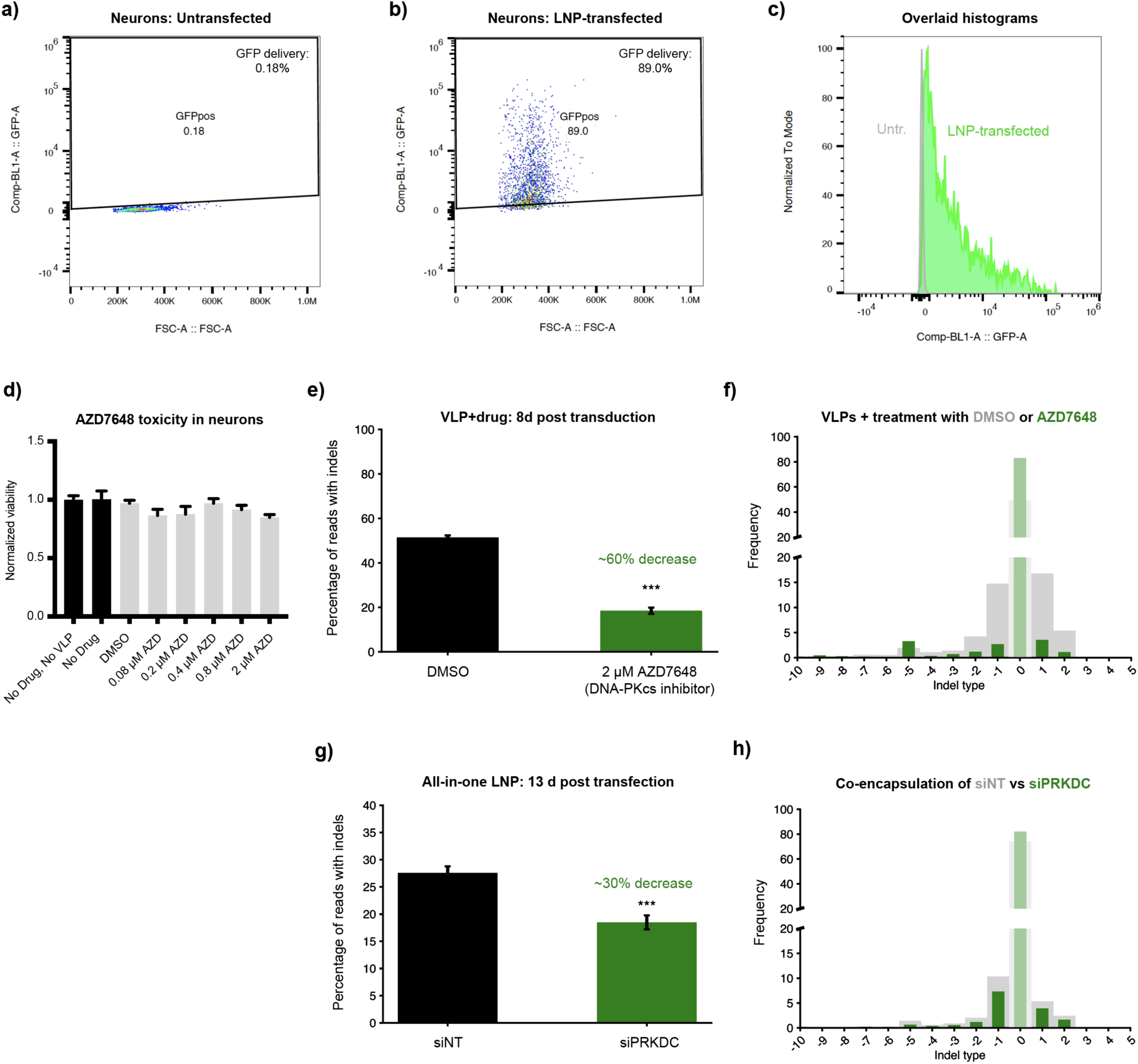
Lipid nanoparticles allow all-in-one delivery of Cas9, sgRNA, and siRNAs to manipulate editing outcomes. **a-c)** Lipid nanoparticles transfect neurons with almost 90% efficiency. Neurons were transfected with LNPs encapsulating GFP mRNA at Day 17+ of differentiation. GFP fluorescence was measured by flow cytometry one week later. **d)** Toxicity of escalating doses of DNA-PKcs inhibitor AZD7648 in neurons, alongside Cas9-VLP treatment. Maximum tested dose was 2 µM, and still tolerable (arbitrary viability threshold of 0.75 as per Extended Data Figure S13). PrestoBlue viability assay at 8 days post-transduction, normalized to age-matched untreated neurons on the same plate. **e-f)** DNA-PKcs inhibitor AZD7648 reduces indels overall in B2Mg1 VLP-treated neurons. 6 replicate wells per condition, treated in parallel. For e: One-Factor ANOVA, *** p<0.0005. **g-h)** All-in-one LNPs co-encapsulating siRNAs against *PRKDC* reduce indels overall in B2Mg1 edited neurons. The effect of *PRKDC* RNA-level inhibition on all-in-one LNP-mediated editing mirrors the effect of *PRKDC* protein-level inhibition on VLP-mediated editing. 6 replicate wells per condition, treated in parallel. For g: One-Factor ANOVA, *** p<0.0005.

## METHODS

### iPSC maintenance

iPSCs were cultured on matrigel-coated 10 cm plates at 37 °C, 85% humidity, and 5% CO2. iPSCs were fed with mTeSR Plus media (StemCell Tech #100-0276) every other day. Optionally, if fed with double the feeding volume of mTeSR Plus one day after passaging, iPSC media could be left unchanged for two days. Upon reaching 80% confluence, iPSCs were passaged 1:10 or 1:20 and treated with 10 μM ROCK inhibitor (Y-27632 dihydrochloride, eg Tocris #1254). For maintenance, ReLeSR (StemCell Tech #100-0483) was used to passage iPSCs as small colonies roughly twice per week. For seeding specific numbers of cells for experiments, Accutase (StemCell Tech #07920) was used to replate iPSCs as single cells after counting.

Cell lines were routinely verified as mycoplasma negative throughout the study. Cell lines used: WTC-NGN2 was used for all experiments except Fig. 3, which used WTC-NGN2-CRISPRi. WTC-NGN2 is the WTC11 iPSC line (Coriell GM25256) with the dox-inducible NGN2 differentiation cassette integrated in the AAVS1 locus.

### Neuron differentiation

Neurons were derived from WTC-NGN2 iPSCs following a differentiation protocol adapted from Tian et al, Neuron (2019), PMID: 31422865. Note however that instead of naming the first day of differentiation Day -3, we name it Day 0. Refer to **Supplemental Table 2** for our adapted differentiation protocol and spreadsheet to aid in calculations.

On Day 3 of differentiation, neurons were seeded onto PDL-coated culture plates (eg Corning #354640, #356414, #356413, #354469): 96-well plates for editing assays, 24-well plates for flow cytometry assays, 6-well plates for RNA assays, or 10 cm plates for ChIP-qPCR. Critically, to maintain neuron viability and reduce media evaporation, we added PBS to the unused wells surrounding cell-seeded wells, especially in 96-well plates. Additionally, to reduce neuronal peeling, for media changes from Day 10 onward we typically removed only half of the existing media volume per well and added a full feeding volume – except when adding VLPs/LNPs/drugs, for which full media changes were used to accurately control concentrations.

In 96-well plate format, each well contained ∼20,000 cells and was treated with 100 μL of VLP- or LNP- containing media. In larger plate formats, these ratios were scaled up proportionally. Note: to transduce 20,000 iPSCs on the same day as the neurons, 10,000 iPSCs were seeded per well one day prior, or 5,000 iPSCs per well were seeded two days prior. Whereas for neurons, 20,000 were seeded on Day 3 of differentiation.

### VLP production and transduction

For HIV-based VLPs (also known as enveloped delivery vehicles or EDVs), we followed the protocols previously described in Hamilton et al, Nat Biotechnol (2024), PMID: 38212493. For FMLV-based VLPs, refer to **Supplemental Table 3** for our full production protocol and calculations.

For both particle types, each “batch” of VLPs consisted of six 10 cm dishes of transfected HEK 293FTs. 44-48 hours post-transfection, each batch’s supernatant was harvested, purified using Lenti-X Concentrator (Takara #631231), and concentrated into 200 μL of OptiMEM (eg Gibco #31985062). Dosage: VLP doses listed in figure captions (either 1 μL or 2 μL as specified) refer to how many μL of this concentrated VLP solution were added per 100 μL of cell culture media, for transduction.

For DSB imaging experiments, transduction was done at Day 14 of differentiation. For all other experiments, transduction was done at Day 17+.

### LNP production and transfection

Lipid mixtures for LNPs were prepared according to previously published procedures (https://doi.org/10.1021/acs.biochem.3c00371). Briefly, stock solutions (10 mg/mL) of MC3 (MedKoo, cat. # 555308), DOPE (Avanti Polar Lipids, cat. # 850725), cholesterol (Sigma-Aldrich, cat. # C8667), and DMG-PEG (Avanti Polar Lipids, cat. # 880151) were individually dissolved in ethanol, while GL67 (N4-Cholesteryl-Spermine HCl Salt, Avanti Polar Lipids, cat. #890893) was dissolved in DMSO. These lipid stock solutions were stored at −30 °C until use. Prior to LNP formation, the lipid solutions were thawed on ice and vortexed as needed. The cholesterol solution was warmed at 40-50 °C to dissolve any crystals that formed during cold storage. Subsequently, MC3, DOPE, cholesterol, DMG-PEG, and GL67 lipids were mixed in molar ratios of 30.8:20.8:32.2:1.2:15, respectively. RNA (1 μg/μL) was dissolved in 200 mM citrate buffer (pH 4), aliquotted, and stored at -80 °C.

Shortly before use, RNA was diluted to 375 ng/μL, then combined with the lipid solution at a 3:1 volume ratio of aqueous phase to lipids. The resulting LNP mRNA complexes were gently vortexed or triturated, and incubated at room temperature for 5-10 minutes. Finally, the LNPs were mixed with the appropriate volume of cell culture media, and added to cells during a full media change. Dosage: we added 300 μL of cell culture media per 4 μL of LNP solution (1 μL of which is lipids). This resulted in an RNA dosage of 125 ng total RNA per 100 μL cell culture media.

Chemically modified GFP mRNA (cat. # L-7201) and Cas9 mRNA (cat. # L-7206) were purchased from TriLink. Chemically modified sgRNAs were ordered from IDT; refer to **Supplemental Table 4** for ordering instructions and resuspension instructions. For siRNAs, TriFECTa DsiRNA kits were ordered from IDT; we used the default TriFECTa kit targeting each gene of interest (cat. #s: hs.Ri.PRKDC.13, hs.Ri.RRM1.13, hs.Ri.RRM2.13, hs.Ri.POLL.13, hs.Ri.XRCC5.13, hs.Ri.XRCC6.13, hs.Ri.DCLRE1C.13, hs.Ri.FANCM.13, hs.Ri.FANCD2.13).

When delivering Cas9 mRNA and sgRNA, total RNA mass inside the particle was split 1:1 between Cas9 mRNA and sgRNA. For all-in-one particles co-delivering siRNAs as well, siRNAs were included at appropriate concentration to yield 1 nM (for Fig. S16g-h), 5 nM (for Fig. 4c-d), or 10 nM (for Fig. 4e) final concentration of “siRNA mixture” in wells. The amount of Cas9 mRNA + sgRNA was reduced proportionally in each case to keep the total RNA concentration the same. Each “siRNA mixture” is a 1:1:1 mixture of the 3 individual siRNAs contained in the IDT TriFECTa kit for a given gene of interest – or, for siNT, it is the TriFECTa kit’s included non-targeting negative control siRNA (labeled NC-1), at the equivalent concentration of total siRNA.

For all LNP experiments, transduction was done at Day 17+ of differentiation.

### Genomic DNA extraction, NGS, and editing analysis

All gDNA for editing experiments was harvested using QuickExtract (#QE09050). After removing cell culture media, 25 μL of QuickExtract was dispensed into each well, and cells were scraped and collected into PCR tube strips (eg Genesee #27-125) or tear-away PCR plates (4titude #4ti-0750/TA). Samples were incubated in a thermocycler for 65 °C for 20 mins, then 98 °C for 20 mins. Extracted gDNA was then stored at -20 °C for short term storage or -80 °C for long term storage.

PCR amplification was done with NEB Q5 master mix (NEB #M0492), and 34 cycles of amplification. Amplicons were then purified using PCR cleanup beads from the UC Berkeley DNA Sequencing Facility, with at least 15 minutes of post-ethanol drying time, and eluted in 30-40 μL of DEPC-treated water. Finally, purified samples were submitted to the UC Berkeley / IGI NGS Core for sequencing via Illumina iSeq (2x150), with 20,000 reads per sample. We processed the resulting sequencing files in Geneious Prime, then used CRISPResso2 (DOI:10.1038/s41587-019-0032-3. PMID: 30809026) to analyze editing outcomes.

For sgRNA spacer sequences and amplicon-NGS primers, refer to **Supplemental Table 5**.

### Drug treatments

Small molecules were resuspended as advised by the manufacturers. Stock concentrations were then prepared at 1000x the desired concentration: refer to **Supplemental Table 6** for stock and final concentrations, as well as catalog numbers. Desired final concentrations were determined by measuring neuronal viability (via PrestoBlue) after escalating drug doses, as shown in **Extended Data Figures S13 and S16**. Per manufacturer suggestions, gemcitabine was resuspended in cell culture grade water; other drugs were resuspended in DMSO.

Drug treatment during Cas9-VLP editing experiments was begun one day prior to transduction. To reduce neuronal peeling from excessive media changes, we did not remove any media during the one-day pretreatment. Instead, we added one feeding volume on top of the existing media, with double the desired drug concentration, to achieve the desired final concentration in the well. The following day (the day of transduction), we performed a full media change, adding media mixed with the desired final concentration of drug and desired volume of VLPs.

### PrestoBlue viability assay

We performed a full media change on neurons, adding in 10% PrestoBlue Cell Viability Reagent (Invitrogen #A13261) by volume, and incubated the cells at 37 °C for 1-2 hours prior to analysis. In 3 control wells with no cells, media with 10% PrestoBlue reagent was added to gauge background fluorescence. After this incubation, the plate was read on a Molecular Devices SpectraMax plate reader and analyzed using the SoftMax software. The average background fluorescence from the control wells was subtracted from all experimental values.

### DSB marker staining and imaging

To stain and image markers of DSB repair (γH2AX and 53BP1), neurons were first fixed with 4% PFA for 15 minutes at room temperature (RT), washed with PBS, and permeabilized with 0.5% Triton X-100 in PBS for 5 minutes at RT. Neurons were then washed with PBS, and incubated with blocking buffer (1% BSA and 0.1% Triton X-100 in PBS) for 1 hour at RT. Then, neurons were incubated with the following buffers at RT, with two PBS washes after each incubation: primary antibody solution for 1 hour (Mouse Anti-phospho-Histone H2A.X Ser139 Antibody, clone JBW301, Sigma #05-636, 1:4000 diluted in blocking buffer; Rabbit Anti-53BP1 Antibody, Novus #100-305, 1:1000 diluted), secondary antibody solution for 1 hour (Goat anti-Mouse IgG H+L 568, Invitrogen #A-11031; Goat anti-Rabbit IgG H+L 488, Invitrogen #A-11034; both 1:1000 diluted in blocking buffer), then DAPI for 2 minutes (1:1000 diluted in PBS, Thermo #62248). Finally, PBS was added to each well, and plates were stored foiled at 4 °C until ready to image.

Higher magnification DSB imaging was performed on a Nikon spinning disk confocal microscope, equipped with a Yokogawa CSU-W1 spinning disk unit, a 60x oil immersion objective lens (N.A. 1.49), Photometrics BSI sCMOS camera and Tokai Hit stage top incubator to maintain temperature, CO2 and humidity. Lower magnification DSB imaging was performed on a BioTek Lionheart LX Automated Microscope, using 40x magnification. For these DSB imaging experiments, neurons were cultured in Ibidi chamber slides (eg Ibidi #80826) with 300 μL of feeding volume.

### Neuronal purity staining and imaging

Neurons were fixed with 4% PFA, washed twice with PBS, then incubated with blocking buffer at RT for 30-60 minutes (5% normal goat serum and 0.1% triton in PBS). After removing blocking buffer and washing twice with PBS, primary antibody solution was added (desired primary antibodies diluted appropriately in PBS with 3% normal goat serum), for a 1 hour incubation at RT. After removing this solution and washing 3 times with PBS, secondary antibody solution was added (appropriate secondary antibodies diluted 1:500 in PBS with 3% normal goat serum, along with 1:1000 diluted DAPI), for a 1 hour incubation at RT in the dark. Following 3 more PBS washes, PBS was added to the wells, and plates were stored foiled at 4 °C until ready to image.

Primary antibodies and their respective dilutions: rabbit anti-Ki67 (1:100, Abcam #ab16667), rat anti-NeuN (1:500, Abcam #ab279297), rabbit anti-TUBB3/Tuj1 (1:500, Sigma #T2200). Secondary antibodies, all used at 1:500 dilutions: goat anti-rabbit 488 (for DAPI-Ki67 combination, Invitrogen #A11008), goat anti-rat 488 (for DAPI-NeuN-TUBB3 combination, LifeTech #A11006), goat anti-rabbit 647 (for DAPI-NeuN-TUBB3 combination, Invitrogen #A21245). These imaging experiments were performed on a CellInsight CX7 microscope, with neurons cultured in PDL coated black/clear 96-well plates (eg Corning # 354640). Images were analyzed by HCS Studio SpotDetector (Ki67 analysis) and CellProfiler (NeuN analysis).

### Flow cytometry

To dissociate neurons for flow cytometry, culture media was removed and then neurons were washed gently with PBS. Papain (reconstituted to 20U/mL in PBS, Worthington #LK003178) was added and incubated for 10 minutes at 37C: 500 or 125 μL papain per well of a 6- or 24-well plate, respectively. Papain was then quenched with DMEM (Corning #10-013-CV) with 10% FBS (eg Avantor #1500-500 or Cytiva #SH30071.03) at 3-5x the papain volume, and pipetted around the edges to lift and collect the sheet of neurons. Neurons were then pelleted, resuspended in 100-500 μL of PBS per sample, and triturated gently to singularize. These samples were passed through strainer-capped FACS tubes (eg Stellar Sci #FSC-9005), and analyzed on an Attune NxT flow cytometer. Results were interpreted using FlowJo.

### Bulk RNA-seq

RNA-seq was performed on 3 replicate samples from each condition, for a total of 24 samples overall: neurons and iPSCs, each transduced with B2Mg1/NEFLg1/NTg1 VLPs (HIV) or untransduced. Cells were transduced in 6-well plate format with 500,000 cells per well, using 25 µL of HIV VLP in 2 mL of media per well. This dose corresponds to 1 µL of VLP per 20,000 cells, or 1.25 µL of VLP per 100 µL media.

Harvest timepoints for each cell type were selected based on their respective time courses of indel accumulation, per Fig. 2. The chosen timepoint for each cell type corresponds to when some, but not all, of the editing has occurred – so that DSB repair is actively ongoing at the time of harvest. This timepoint is 3 days post-transduction for neuron samples, and 1 day post-transduction for iPSC samples, which also avoids confounding effects from cell proliferation and/or dilution. Untransduced cells were harvested on the same day as the transduced cells.

RNA was extracted using the Quick-RNA™ Microprep Kit (cat. #R1050). Using 500 ng of total RNA, we prepared the mRNA libraries using the QuantSeq 3’ mRNA-Sequencing Library FWD V1 Prep Kit (cat. #015.96). After cDNA synthesis, we used 17 PCR cycles to amplify the libraries. Following bead purification, mRNA concentrations were determined by Qubit and fragment lengths were quantified using High Sensitivity d5000 Reagents (cat. #5067-5593) on the Agilent TapeStation 4200. We normalized our libraries to 8.25 nM for pooling and sequenced through single-end sequencing on the Illumina NovaSeqX 10B flow cell with read lengths 101x12x24 (Read 1, Index 1, Index 2). Sequencing was performed at the UCSF Center for Advanced Technologies (CAT).

Sequencing reads were trimmed using CutAdapt (DOI:10.14806/ej.17.1.200) and aligned using HISAT2 (DOI:10.1038/s41587-019-0201-4), and then a read count matrix was generated using featureCounts (DOI: 10.1093/bioinformatics/btt656). Differential expression analysis was performed on this count matrix using EdgeR (DOI:10.1093/bioinformatics/btp616). Functions within EdgeR used to statistically determine differentially expressed genes were: glmQLFit, glmQLFTest (with FDR for adjusted p-values), and decideTestsDGE. For venn diagrams and enrichment analysis, additional tools used were topTags with Benjamini-Hochberg correction for adjusted p-values, and WebGestalt (https://www.webgestalt.org/#).

### ChIP-qPCR

For each timepoint condition (untransduced, 2 day, 4 day, 8 day), 20 million neurons were grown across 2 10 cm dishes per condition (10 million neurons per plate). All 8 of these plates were cultured in parallel, during the same batch of differentiation. At Day 17, all transduced plates were transduced with 2 μL of B2Mg1 VLP (FMLV) per 100 μL media (total of 200 μL VLP per plate). Untransduced plates were harvested at Day 17. Remaining plates were harvested 2/4/8 days post-transduction as labeled.

At each harvest timepoint, 2 10 cm dishes were fixed in parallel: one for each ChIP pulldown (Mre11 and γH2AX). Cells were fixed, pelleted, and snap frozen per ActiveMotif’s ChIP fixation protocol (https://www.activemotif.com/documents/1848.pdf), then submitted to ActiveMotif for ChIP-qPCR. ChIP-qPCR was performed using the antibodies and qPCR primers listed in **Supplemental Table 7**.

## DATA AVAILABILITY

Raw and processed RNA sequencing files are available through the NCBI Gene Expression Omnibus (GEO), via accession code GSE272812: https://www.ncbi.nlm.nih.gov/geo/query/acc.cgi?acc=GSE272812.

## ETHICS STATEMENT

Studies in the Conklin Lab involving human induced pluripotent stem cells were reviewed and approved by the UCSF Institutional Review Board. The donor from whom the WTC iPSC line was derived provided written informed consent for the generation and use of their iPSCs, which are commercially available through Coriell (GM25256).

## AUTHOR CONTRIBUTIONS

GNR and BRC conceived the study. BRC, MK, and GNR oversaw the overall project directions and planning. GNR, SJN, KGC, MS, and BLM performed most experiments. JRH, CF, BSP, and CRSE designed and/or produced VLPs. RS and NM designed and/or produced LNP formulations. J-CL imaged DSB foci. JJ performed RNAseq library prep. MPM and SHL contributed to supplementary data. HLW, LMJ, AN, BA, and JAD provided conceptual and critical guidance, and helped shape the manuscript. GNR wrote the manuscript with input from all authors.

## ACKNOWLEDGMENTS

We thank Zachary Nevin, Beeke Wienert, and all other members of the Conklin Lab for their helpful suggestions and feedback. We are also grateful to Jacob Corn, Alexis Komor, Jeffrey Hussmann, John Doench, Françoise Chanut, Netravathi Krishnappa, Michael Ward, Andy Qi, Hesong Han, Abdullah Syed, Bjoern Schwer, James Dahlman, Zoe Grant, David Toczyski, and Tippi Mackenzie for their advice. This work was enabled by the Gladstone Stem Cell, Flow Cytometry, Assay Development & Drug Discovery, and Bioinformatics Cores, as well as ActiveMotif, the IGI Next Generation Sequencing Core, and the UCSF Center for Advanced Technology.

